# Decoding the porcine developing spatial processing system and production of human entorhinal stellate cell-like cells by a direct programming approach

**DOI:** 10.1101/738443

**Authors:** Tobias Bergmann, Yong Liu, Leo Mogus, Julie Lee, Ulrich Pfisterer, Louis-Francois Handfield, Andrea Asenjo-Martinez, Irene Lisa-Vargas, Stefan E Seemann, Jimmy Tsz Hang Lee, Nikolaos Patikas, Birgitte Rahbek Kornum, Mark Denham, Poul Hyttel, Menno P Witter, Jan Gorodkin, Tune H Pers, Martin Hemberg, Konstantin Khodosevich, Vanessa Jane Hall

## Abstract

Classic studies investigating how and when the entorhinal cortex (component of the memory processing system of the brain) develops have been based on traditional thymidine autoradiography and histological techniques. In this study, we take advantage of modern technologies to trace at a high resolution, the cellular complexity of the developing porcine medial entorhinal cortex by using single-cell profiling. The postnatal medial entorhinal cortex comprises 4 interneuron, 3 pyramidal neuron and 2 stellate cell populations which emerge from intermediate progenitor and immature neuron populations. We discover four MGE-derived interneurons and one CGE-derived interneuron population as well as several IN progenitors. We also identify two oligodendrocyte progenitor populations and three populations of oligodendrocytes. We perform a proof-of-concept experiment demonstrating that porcine scRNA-seq data can be used to develop novel protocols for producing human entorhinal cells in-vitro. We identified six transcription factors (*RUNX1A1, SOX5, FOXP1, MEF2C, TCF4, EYA2*) important in neurodevelopment and differentiation from one *RELN*+ stellate cell population. Using a lentiviral vector approach, we reprogrammed human induced pluripotent stem cells into stellate cell-like cells which expressed *RELN, SATB2, LEF1* and BCL11B. Our findings contribute to the understanding of the formation of the brain’s cognitive memory and spatial processing system and provides proof-of-concept for the production of entorhinal cells from human pluripotent stem cells in-vitro.

## Introduction

Uncovering the temporal events in the emergence of different cell types in the developing entorhinal cortex (EC) is paramount for understanding the formation of cognitive memory and spatial processing. The EC has a pertinent role in the processing of memory and navigation, which is dependent on its intrinsic organization and extrinsic connectivity. Specific navigation-related functions have been discovered in the EC including object recognition (Ridley, Samson, Baker, & Johnson, 1988), processing speed movement (Kropff, Carmichael, Moser, & Moser, 2015), processing location (Quirk, Muller, Kubie, & Ranck, 1992), processing head-direction (Taube, Muller, & Ranck, 1990), recognizing proximal borders (Solstad, Boccara, Kropff, Moser, & Moser, 2008) and processing working memory (Olton, Walker, & Wolf, 1982). These functions signify the importance of the EC in spatial navigation and arise from subsets of firing cells from either the medial EC (MEC) or lateral EC (LEC). Extensive anatomical and neuronal electrophysiological classification of the EC has been performed (Canto & Witter, 2012a, 2012b; Canto, Wouterlood, & Witter, 2008) which has recently been supplemented with molecular characterization by single-cell RNA sequencing (scRNA-seq) in the adult EC (Grubman et al., 2019; Leng et al., 2021). Currently, none, or only very few genes can be used to identify the phenotypically identified cells, such as the grid cells, head-direction cells and speed cells and the locations of these cell types are only partially understood. In some cases, these phenotypic attributes appear to be derived from anatomically different cell types. For example, within the MEC, the grid cells are located in both the dorsal and ventral MEC (Stensola & Moser, 2016) and found across all cortical layers, though showing a preference in layer (L) II (Sargolini et al., 2006). However, the grid cells are considered to represent principal neurons including both stellate cells (SCs) and pyramidal neurons (Domnisoru, Kinkhabwala, & Tank, 2013; Rowland et al., 2018; Schmidt-Hieber & Hausser, 2013) suggestive of networking interrelations that govern their function. Gene markers that distinguish the SCs from the pyramidal neurons are Reelin (*RELN*) and Calbindin (*CALB1*), respectively (Fuchs et al., 2016; Leitner et al., 2016; Perez-Garcia et al., 2001; Seress, Leranth, & Frotscher, 1994; Varga, Lee, & Soltesz, 2010; Witter, Doan, Jacobsen, Nilssen, & Ohara, 2017), although a subpopulation of both types expresses both markers (Witter et al., 2017). Reelin is also expressed in Fan cells which reside in the superficial layer of the LEC and these neurons are important for object recognition (Germroth, Schwerdtfeger, & Buhl, 1989; Nilssen et al., 2018). A specialized grid cell, the conjunctive grid cell, fires when a grid vertex is crossed upon heading in a specific direction. It is absent from LII, mostly present in LIII and LIV, but can also be found in LVI (Sargolini et al., 2006). The head-direction cells are principal neurons found in LII to LVI and are absent from LII (Giocomo et al., 2014; Sargolini et al., 2006). The border cells are also considered to be principal neurons and are found within all the MEC layers (Solstad et al., 2008) which often overlap with a SC identity (Tang et al., 2014). The speed cells map to GABAergic inhibitory neurons which express parvalbumin (PVALB), but not somatostatin (SST) (Ye, Witter, Moser, & Moser, 2018).

A clearer understanding of the molecular landscape of the EC may help to reconcile the anatomical, functional and connective profiles of the cell types that exist in the EC which would further improve our understanding of spatial navigation processing. One single-cell RNA sequencing (scRNA-seq) study was recently published on the EC from Alzheimer’s disease (AD) patient and healthy brains, however, the vast majority of cells captured in their dataset were oligodendrocytes (Grubman et al., 2019). A second study has highlighted the transcriptional profiles of several excitatory neurons and glia in the adult EC, but does not decipher the cell types residing in the LEC or the MEC (Leng et al., 2021). Further, in both studies the research concentrated on differences between healthy and disease state rather than on the characterization of the cell types *per se*. We envisage a more comprehensive analysis of the EC cell types as they emerge during development can provide further insight into the formation processes within the EC and insight into transcriptional changes that regulate maturation and connectivity of the EC.

Studying the signaling cascades that regulate the differentiation of brain cell types has resulted in novel protocols for producing several neuronal cell subtypes in-vitro from pluripotent stem cells. Protocols exist for production of, for example, multipotent neuroepithelia (Falk et al., 2012; W. Li et al., 2011; Nat et al., 2007), radial glia (Bamba et al., 2014; Bouhon, Joannides, Kato, Chandran, & Allen, 2006; Duan, Peng, Pan, & Kessler, 2015; Gorris et al., 2015; Hofrichter et al., 2017; Liour et al., 2006; Nat et al., 2007; Onorati et al., 2010; Reinchisi, Limaye, Singh, Antic, & Zecevic, 2013), glutamatergic neurons (Cao et al., 2017; Chuang, Tung, Lee-Chen, Yin, & Lin, 2011; Reyes et al., 2008; Thoma et al., 2012), GABAergic neurons (Barretto et al., 2020; Shin, Palmer, Li, & Fricker, 2012; Sun et al., 2016; Yang et al., 2017), midbrain dopaminergic neurons (Daadi, 2019; Jo et al., 2016; Jovanovic et al., 2018; Kikuchi et al., 2017; Kriks et al., 2011; Studer, 2012; Tofoli et al., 2019; Xue et al., 2019; P. Zhang, Xia, & Reijo Pera, 2014), 5-HT neurons (Bethea, Reddy, Pedersen, & Tokuyama, 2009; Kumar, Kaushalya, Gressens, Maiti, & Mani, 2009; Lu et al., 2016; Shimada et al., 2012; Vadodaria, Marchetto, Mertens, & Gage, 2016), motor neurons (Bianchi et al., 2018; Chipman, Toma, & Rafuse, 2012; Dimos et al., 2008; Sances et al., 2016) and enteric neurons (Barber, Studer, & Fattahi, 2019; Workman et al., 2017). This process can be orchestrated via indirect differentiation using exogenous factors and/or small molecules added to the media, or by directly programming the cells into the desired end phenotype by using targeted overexpression of regulatory transcription factors (K. Li et al., 2016; Oh & Jang, 2019; Qin, Zhao, & Fu, 2017). Producing neurons of the EC from human iPSCs would provide a useful model for studying the functions of individual cell types but also be useful in modeling diseases which affect the EC, such as AD. Recent research has highlighted that both Reelin+ and *RORB*+ neurons in the EC are particularly vulnerable to either accumulating Amyloid beta, neurofibrillary tangles or dying early in the disease (Kobro-Flatmoen, Nagelhus, & Witter, 2016; Leng et al., 2021) and studying these cell types further will help to elucidate why select populations of neurons in the EC are particularly vulnerable to the disease. We therefore aim to determine the key regulatory transcription factors in entorhinal Reelin+ cells in order to create a novel direct programming protocol for producing these neurons in-vitro. In light of difficulties in obtaining human fetal tissues from the second and third trimester to investigate EC development, we used an alternate large mammal, the pig. The pig develops a gyrencephalic brain midway through gestation, similar to humans and our recent research confirms it to be an excellent model of the developing human EC that recapitulates neurogenesis more closely in the human than rodents (Liu et al., 2021). We focus our attention on the MEC given the phenotypes and connectivity of several cell types are well studied but the molecular genotypes residing within the MEC are not well delineated.

In this study, we determine the molecular profiles of the neural cell types within the developing and postnatal MEC using scRNA-seq and identify entorhinal specific neuron subtypes. We then identify potential regulators of differentiation in interesting *RELN*+ excitatory neuron populations by performing differential expression of transcription factors. Focusing on one *RELN*+ population, we transduce six selected transcription factors using a lentiviral overexpression approach and directly program human induced pluripotent stem cells (iPSCs) into *RELN*+ entorhinal SC-like progenitors.

## Results

### ScRNA-seq reveals thirty two cell populations in the developing MEC

ScRNA-seq is a powerful approach for revealing the molecular identity of cell types in the different types of tissue, including the brain. We used this methodology to investigate the timing in the emergence of different cell types of the EC and their molecular identity during development and postnatal maturation. ScRNA-seq was performed using the 10X Genomics microfluidics-based Chromium platform on whole-cells isolated from the EC at embryonal day 50 (E50), E60, E70 and postnatal day 75 (P75), and isolated nuclei in the later stages of development at E70 and P75 to ensure efficient capture of the neurons in the tissue (Figure 1A). We chose to integrate datasets for all age groups into a one-dimension reduction analysis as opposed to analyzing time points separately to ensure superior normalization and to allow us to easily cross-label cell types that existed both during fetal development and in the postnatal brain. We excluded red blood cells (expressing hemoglobin subunits *HBB, HBE1, HBM, HBZ* and on average, only 291 genes per cell) and vascular cells (*PDGFRB*^+^*/PECAM1*^+^) from our dataset, which left 24,294 cells (mean of 2798 different genes per cell) for further analyses. We log-normalized each sample dataset, merged all samples, batch corrected by FastMNN (Haghverdi, Lun, Morgan, & Marioni, 2018) and performed cell clustering in Seurat (Figure 1A). Clustering of the merged dataset revealed thirty-two distinct cell clusters (Figure 1B) with many clusters represented by cells from datasets across different batches with good integration of batched within same age (Figure 1C). The clusters could be delineated into 6 main cell populations demarcated by canonical markers for intermediate progenitor cells (IPs, *PAX6*^+^*/EOMES*^+^*/NEUROD1*^+^), excitatory neurons (Excitatory, *TBR1*^+^*/EMX1*^+^*/SATB2*^+^*/BCL11B*^+^*/SLC17A6*^+^*/SLC17A7*^+^*/DCX*^+^), GABAergic interneurons (IN,*GAD1*^+^/*GAD2*^+^/*DCX*^+^), microglia (Microglia, *AIF1*^+^*/CXCR3*^+^*/PTPRC*^+^*/CSF1R*^+^), oligodendroglia (Oligo, *MBP*^+^*/CLDN11*^+^*/OLIG1*^+^*/PDGFRA*^+^), astrocytes (*AQP4*^+^*/GFAP*^+^*/GLAST*^+^) and radial glia (RGC, *HES1*^+^*SOX2*^+^) (Figure 1D). Cluster 30, contained a small number of cells (n=70) which contained twice as many transcripts and genes expressed (Figure 1C) which we believed to be doublets and which failed to be removed by the data-preprocessing. Cluster 31 expressed *MBP* (Figure 1C) but was also omitted from the subanalysis, as the highest enriched genes in this cluster were mitochondrial genes (Figure S1A) and may likely be dying cells. Furthermore, we validated our marker-expression driven annotations by projecting our dataset using scmap (Kiselev, Yiu, & Hemberg, 2018) onto a publicly available and annotated datasets of the human embryonic prefrontal cortex (Zhong et al., 2018) and the human adult middle temporal gyrus (Hodge et al., 2019) (Figure 1E). We found a high concordance between our annotation and the human brain datasets, despite the obvious regional differences.

**Figure 1.**
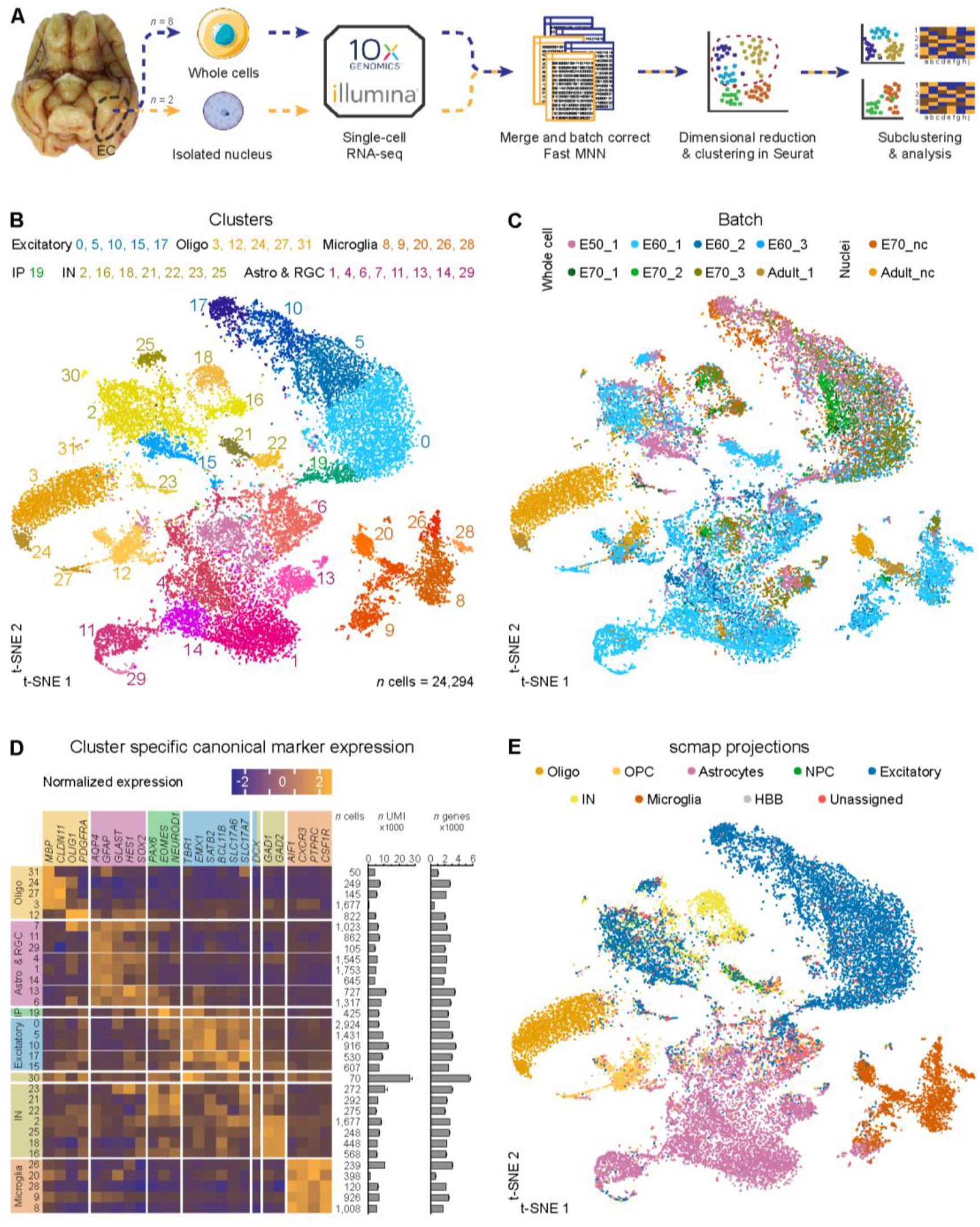
Single-cell profiling of the entorhinal cortex reveals thirty-two distinct cell clusters of major cell type populations. **(A)** The computational pipeline includes ten batches of single-cell and single-nuclei isolated cells from E50, 255 E60, E70 and P75 which were captured using the 10X genomics platform. Batches were normalized using the Fast MNN approach and dimensional reduction and clustering were performed in Seurat followed by subclustering and analyses of selected clusters. **(B)** A t-SNE plot of 24,294 cells merged from all timepoints revealed thirty-two distinct populations following FastMNN batch correction. Oligodendro-glia/-cytes (Oligo), astrocytes and radial glia (Astro & RGC), intermediate progenitors (IP), excitatory neurons (Excitatory), interneurons (IN) and microglia (Microglia) **(C)** tSNE plot annotated by batch 260 demonstrates good integration of subtypes within same ages following correction by FastMNN of the 10 datasets (pre-batch correction shown in Figure S5C). **(D)** Analysis of canonical marker genes resulted in categorization of several Oligo, Astro & RGC, IP, Excitatory, IN and Microglia clusters. The number (n) of cells, mean UMI counts (unique transcripts) and mean number of different expressed genes for each cluster are also represented. Error bars denote the SD. **(E)** Projection of the dataset onto an already-annotated human fetal prefrontal cortex and human medial temporal gyrus single-cell dataset consolidates 265 marker gene-driven annotation of the dataset. Abbreviations: NPC, neural progenitor cells; HBB, marker expressed in human blood cells.

### Progenitor and adult interneuron (IN) populations derived from the MGE and CGE are detected in the developing MEC

To investigate the temporal emergence of INs within the EC, we sub-clustered a total of 3,498 cells from the dataset using the canonical IN markers, *GAD1* and *GAD2* (clusters 2, 16, 18, 21, 22, 23 and 25) which produced ten subclusters IN0-IN9 (Figure 2A). We included clusters 2 and 25 in our IN analyses even though the scmap projection annotated these two clusters as excitatory neurons (Figure 1D, E). Despite this discrepancy, cluster 25 expressed IN genes, such as *GAD1* and *GAD2* and lacked *SLC17A6/7* and became IN7 after subclustering. For cluster 2 from the main dataset (mainly present at E60), a closer analysis of canonical markers suggested an ambiguous identity as the cells expressed both the excitatory markers, *SLC17A6/7* and IN markers, *GAD1/2* (Figure 1B, D). The UMI counts for this cluster were normal, therefore, it is unlikely to contain doublets. Cluster 2 when subclustered spread across the IN0-IN2 and IN6 populations. Five of the IN populations were only detected during development, suggesting they were IN progenitors (IN1, IN5, IN7, IN8 and IN9). IN1, IN5 and IN7 were detected during earlier embryonic gestation in the pig at E50 and E60, compared to IN8 and IN9 which were detected later, at E70 (Figure 2B). IN5, IN7 and IN8 also clustered further away from the other IN clusters which also suggests they are more transcriptionally distinct than the other IN clusters (Figure 2A). The remaining IN populations, IN0, IN2, IN3 and IN6 were all found in the postnatal brain and are likely adult IN populations (Figure 2B). We could identify several well-known IN markers that were either expressed broadly or label specific IN cell subtypes, including, *CALB1, DLX1, DLX5, GAD1, GAD2, LHX6, NPY, RELN, SST*, and *SOX6*, (Figure 2C). However, we were unable to identify other IN markers, *HTR3A, PVALB, CCK, NOS1, LHX5* and *VIP*, likely due to low expression levels and gene dropout effects.

**Figure 2.**
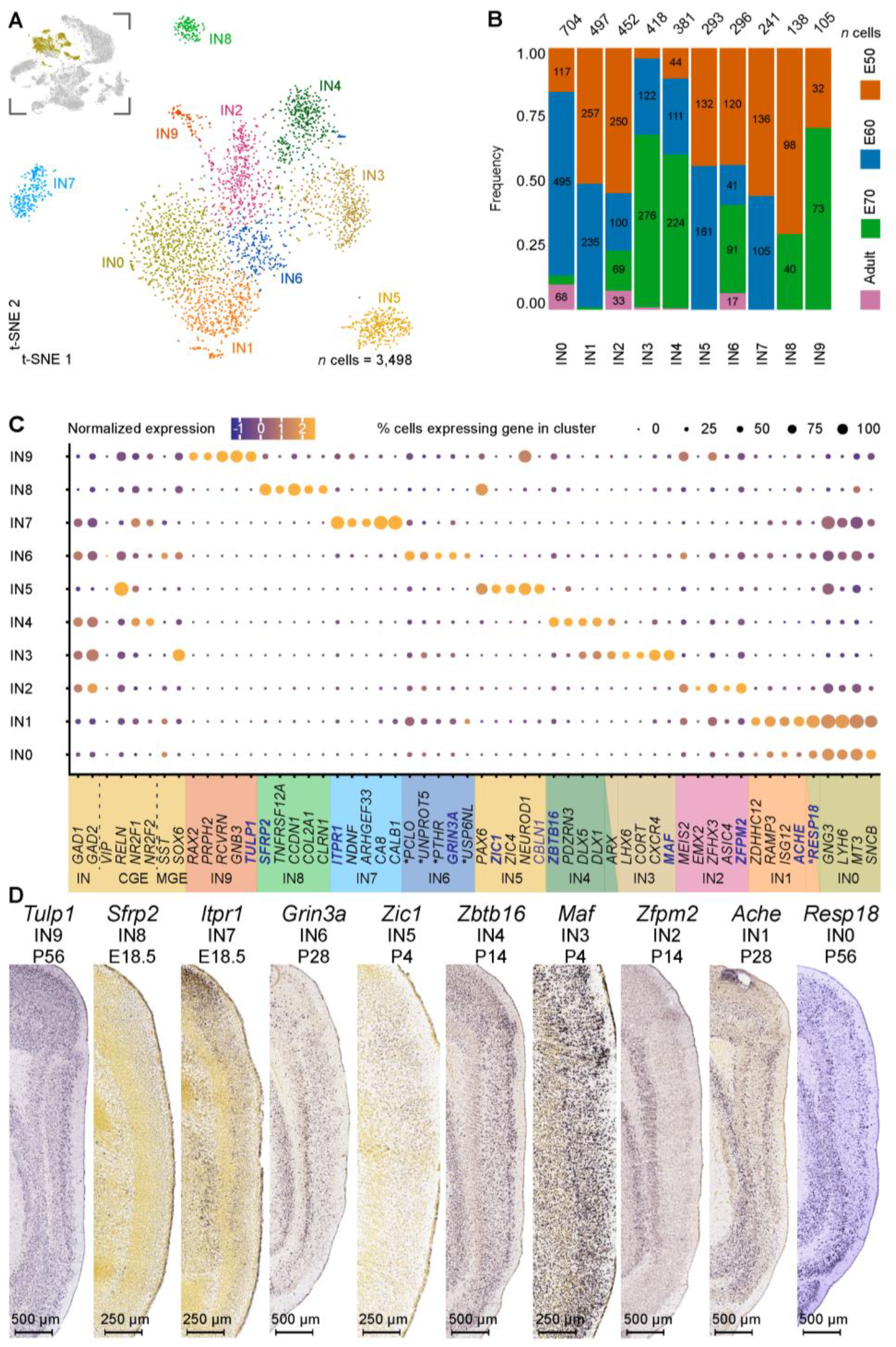
Subclustering of entorhinal cortex (EC) interneurons (INs) reveals ten clusters with unique gene signatures. **(A)** A t-SNE plot of the subclustering identifies ten interneuron (IN) clusters with one cluster more transcriptionally distinct than the others. IN clusters highlighted in yellow in the inset image were subclustered from the parent dataset and are depicted in the top left corner. **(B)** An analysis of the developmental stages across the ten clusters highlights IN1, IN5, IN7, IN8 and IN9 are found exclusively during development and suggestive of IN progenitors and the remaining clusters are adult INs. **(C)** Unique gene signatures were identified for the ten clusters by differential gene expression analysis, together with the population specific expression of canonical markers for interneurons (IN) derived both from the caudal ganglionic eminence (CGE) and the medial ganglionic eminence (MGE). Localization of genes with names in blue are shown in (D). * marks the human ortholog names (Table S1). **(D)** Sagittal sections from the ISH datasets from the Allen Developing Mouse Brain Atlas and Allen Mouse Brain Atlas highlight the location of selected markers from the IN clusters across the EC layers in the developing and postnatal mouse brain (“Allen Developing Mouse Brain Atlas,” 2008; “Allen Mouse Brain Atlas,” 2004; Lein et al., 2007). Image credit: 330 Allen Institute. Scale bars: 250 μm (for E18.5, P4 and P14) and 500 μm (for P56).

Most cortical INs are generated in the ganglionic eminences and are divided in two major classes based on their place of origin: the medial ganglionic eminence (MGE) and caudal ganglionic eminence (CGE) (Gelman & Marin, 2010). The MGE produces PVALB and SST-expressing cortical INs, which account for >60% of all INs in the cortex and the CGE produces a highly diverse group of INs, which in mice express serotonin 5HT3A receptor (Rudy, Fishell, Lee, & Hjerling-Leffler, 2011; Vucurovic et al., 2010) (gene *HTR3A*) However, in human INs, HTR3A labels only a minor population of CGE INs (Hodge et al., 2019), which is also likely the case in pigs, since we did not detect robust *HTR3A* expression in our data. Given we could not identify PVALB in our data, we also used other MGE IN markers including *LHX6* (Liodis et al., 2007) and *SOX6* (Batista-Brito et al., 2009) to identify MGE-derived INs. Based on *SST* expression in IN0, IN2 and *LHX6* and *SOX6* in IN3, we assign these clusters as MGE-derived INs (Figure 2C). It was difficult to assign IN6 to either MGE or CGE INs, given an overlapping expression profile but previous research suggests *GRIN3A*, which was highly expressed in this population, is detected in SST INs (Murillo et al., 2021), suggesting IN6 is likely of MGE origin. The progenitor population IN1 also expressed *SST*. Interestingly, both IN0 and IN1 have relatively similar transcriptional profiles. It is likely that these are unique cell populations since IN0 had lower expression of *RELN* and IN1 expressed higher levels of *RAMP3, ISG12* and *ACHE* and only IN0 could be detected after E70 of development (Figure 2C). However, further phenotypic profiling is required to conclude that these are separate populations. The remaining clusters had more distinct gene signatures. CGE-derived INs were identified based on expression of Coup TF1 (*NR2F1* gene), CoupTF2 (*NR2F2* gene) (Kanatani, Yozu, Tabata, & Nakajima, 2008) and *NDNF (Tasic et al*., *2018*). One population, IN4, expressed *NR2F1* and *NR2F2* and is assigned as a CGE population (Figure 2C). The remaining progenitor populations IN5, IN7-IN9 clusters also expressed these CGE IN markers (Figure 2C).

To investigate the location of the varying IN progenitors and adult populations across the EC layers, we evaluated the expression of unique genes from the clusters in the *in-situ* hybridized EC of the mouse brain from the public databases, the Allen Mouse Brain Atlas (2004) (“Allen Mouse Brain Atlas,” 2004; Lein et al., 2007) and the Allen Developing Mouse Brain Atlas (2008) (“Allen Developing Mouse Brain Atlas,” 2008). IN0 and IN1 both expressed a gene, *RESP18*, which was localized to the superficial layers and deep layers of the adult mouse EC (“Allen Mouse Brain Atlas,” 2004) (Figure 2D), suggesting these INs are not located in the middle cortical layers. In addition *ACHE* was upregulated in IN1 and also found in the superficial and deep layers, confirming the location of at least the IN 1 neurons in the mouse EC (Figure 2D). *ZIC1, ITPR1, SFRP2* and *TULP1* were expressed in the IN progenitor populations IN5, IN7, IN8 and IN9, respectively and were distributed across the layers of the developing and postnatal EC mouse brain. *ZFPM2* was upregulated in IN2 and found only in the deeper layers (Figure 2D). *MAF* and *ZBTB16* were unique upregulated genes in IN3 and IN4, respectively and were expressed across all the EC layers, however, ZBTB16 was also upregulated in the superficial layers suggesting potential high abundance of IN4 CGE INs in the superficial layers. (Figure 2D). *GRIN3A* was uniquely expressed in IN6 and detected across the layers but also upregulated in the deeper layers of the MEC suggesting different localization of these INs between the MEC and LEC (Figure 2D). In sum, we identified five IN progenitor populations (IN1, IN5, IN7-IN9), four adult IN MGE populations (IN0, IN2, IN3, IN6), and one IN CGE population (IN4).

### Single-cell RNA sequencing analysis reveals that OPCs emerge in the early EC

We then assessed the oligodendroglia populations within the EC, which consisted of oligodendrocyte progenitor cells (OPCs) and oligodendrocytes at various stages of maturation. Subclustering and subsequent analyses were performed on clusters 3, 12, 24, 27 (Figure 3A insert). Five distinct clusters were identified from the subclustering (Figure 3A-C). Two clusters (OPC1 and 2) expressed OPC markers *PDGFRA* and *OLIG1* (Figure 3C), with a large proportion of the cells undergoing cell cycle division (50% and 25% respectively, Figure 3D). We were unable to identify *OLIG2* in these two clusters, likely due to low copy numbers and subsequent gene dropout effects. However, this data suggested these two clusters were progenitor cells. Cluster OPC1 was composed of mostly fetal cells from E50 to E70 whilst cluster OPC0 was detected both during gestation and postnatally (Figure 3B). OPC0 and OPC1 are clearly two separate populations, as both populations are detected after birth, albeit the clustering suggests a larger population of OPC0 after birth. Several genes have been previously identified in OPCs. For example, cluster OPC0 expressed the genes *LHFPL3* and *MMP16*, and cluster OPC1, *STMN1;* previously identified in rodent OPCs (Artegiani et al., 2017; Hu et al., 2011; Lin, Mela, Guilfoyle, & Goldman, 2009; Magri et al., 2014). To characterize whether these clusters resided in spatially different niches, we examined their unique gene signatures in the EC from the Allen Mouse Brain Atlas and Allen Developing Mouse Brain Atlas ISH datasets (“Allen Developing Mouse Brain Atlas,” 2008; “Allen Mouse Brain Atlas,” 2004; Lein et al., 2007). We detected *Cntn1* (a marker previously identified in OPCs (Lamprianou, Chatzopoulou, Thomas, Bouyain, & Harroch, 2011)) from OPC0 in the superficial layers in the developing mouse EC (E18.5) and at a later time point across all cell layers, at P28 in the mouse EC (“Allen Developing Mouse Brain Atlas,” 2008; “Allen Mouse Brain Atlas,” 2004) (Figure 3E). We detected the OPC1 marker *Hmgb1* (also previously identified in OPC progenitors (Nicaise et al., 2019)) across all EC cell layers in the postnatal P4 and P28 and adult mouse EC (“Allen Developing Mouse Brain Atlas,” 2008; “Allen Mouse Brain Atlas,” 2004) (Figure 3E) showing the gene as expressed even later, after birth.

**Figure 3.**
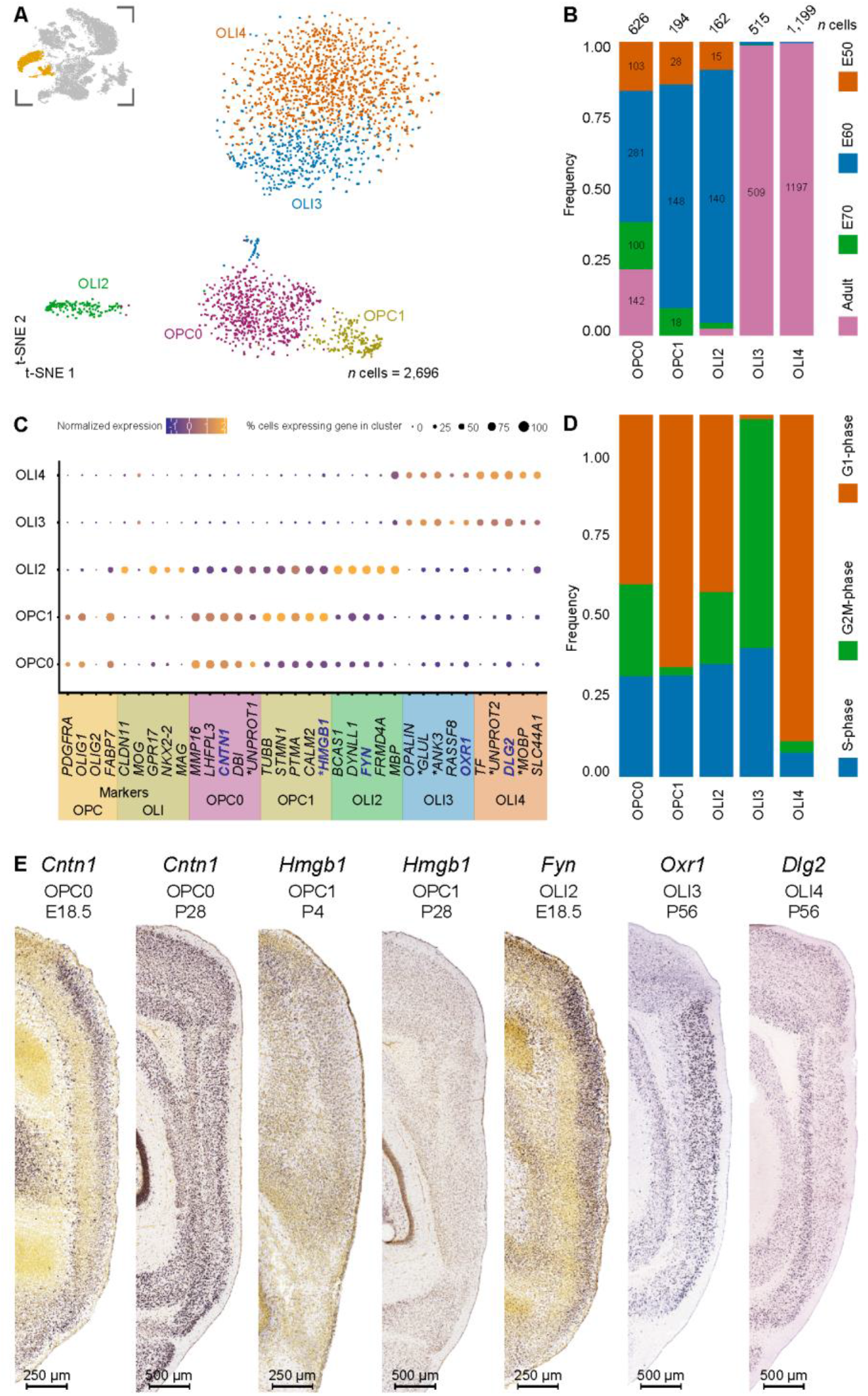
Subclustering of oligodendro-cytes and -glia reveals oligodendrocyte progenitor cells (OPCs) emerge in the early entorhinal cortex. **(A)** Visualization of the oligodendroglia subset (number (n) of cells = 2,696) shows five clusters in a 385 t-SNE plot. Oligodendroglia clusters highlighted in orange in the inset image were subclustered from the parent dataset and are depicted in the top left corner. **(B)** The developmental stages of the oligodendroglia visualized by a bar plot highlight a distinct adult and fetal populations and populations that exist both during development as early as E50 and postnatally. **(C)** Canonical lineage markers and unique gene signatures for the five clusters were identified by differential gene expression analysis. Localization of genes with names in blue is shown in (E). * marks the human ortholog names (Table S1). **(D)** Distribution of 390 cells within different cell cycle phases highlights OLI4 is largely a non-mitotic population and the remaining clusters are mitotically active. Note, G1-phase is indistinguishable from the G0-phase. **(E)** Sagittal sections from the ISH datasets from the Allen Developing Mouse Brain Atlas and Allen Mouse Brain Atlas (“Allen Developing Mouse Brain Atlas,” 2008; “Allen Mouse Brain Atlas,” 2004; Lein et al., 2007) highlight the location of selected markers from the OPC and OLI subtypes. Image credit: Allen Institute. Scale bars: 250 μm (for E18.5 and P4) and 500 μm (for P28 and P56).

Interestingly, our data revealed an additional population, OLI2, that was constituted predominantly of fetal cells from E60 OLI2 (Figure 3A-B). These cells surprisingly also expressed the mature oligodendrocyte markers, *MBP, CLDN11, GRP17, NKX2-2*, and *MAG* (Figure 3C). Together, this data suggests that a small population of myelinating oligodendrocytes might be present in the fetal porcine EC already at E60. The oligodendrocyte clusters OLI3 and OLI4 consisted almost exclusively of adult cells and were highly similar in profile (Figure 3B, C). We speculate these to be mature myelinating oligodendrocytes. Nearly all the OLI3 cells expressed G2M/S-phase cycling genes while OLI4 cells were mostly quiescent and expression genes from the G0/1-phase of the cell cycle (Figure 3D). In the Allen Developing Mouse Brain Atlas, expression of the OLI2 specific marker *Fyn* showed predominant expression in the upper layers of the developing E18.5 mouse EC (“Allen Developing Mouse Brain Atlas,” 2008) (Figure 3E). In the Allen Mouse Brain Atlas, the OLI3 unique gene marker, *Oxr1*, was expressed in LI-III of the adult mouse EC while the OLI4 unique gene marker, *Dlg2*, was found in the deep layers (“Allen Mouse Brain Atlas,” 2004) (Figure 3E-F). In summary, our data indicate the presence of 2 OPC populations that may populate different layers of the EC, with one pre-myelinating oligodendrocyte population and two mature oligodendrocyte profiles with similar profiles.

### Single-cell RNA sequencing reveals unique gene signatures for stellate cells and pyramidal neuron populations

To gain a deeper understanding of the temporal pattern of emerging neurons we performed subclustering on the IP and excitatory neuron populations identified from the parent dataset and uncovered unique gene expression signatures. We subclustered and analyzed a total of 6,833 cells, including the excitatory clusters 0, 5, 10, 15, and 17 and the IP cluster 19. Figure 4A, insert). The subclustering revealed ten distinct populations (Figure 4A). Cells from the gestational stages dominated the subset of IPs and excitatory neurons, with only very few adult cells represented (Figure 4B). This is due to the fact that we captured mostly glia in the postnatal brain samples. Amongst the clusters, one population exhibited a typical IP identity. Cells in cluster IP4 expressed the typical IP markers *EOMES, SOX2*, and *NEUROD4* (Figure 4C) (Pollen et al., 2015) and could be detected across all stages of development, but also in the postnatal brain suggestive of ongoing postnatal neurogenesis (Figure 4B). IP8 was a detected only during the neurogenesis period (E50-E60) (Liu et al., 2021) and expressed typical neural progenitor markers including, *PAX3* (Blake & Ziman, 2014), *DBX1* (Lacin, Zhu, Wilson, & Skeath, 2009) and the ventral midbrain progenitor marker *OTX2* (Puelles et al., 2004). When referring to the Allen Developing Mouse Brain Atlas ISH dataset (“Allen Developing Mouse Brain Atlas,” 2008), the E18.5 mouse EC showed expression of a unique marker *Cxcr4*, across all the layers (Figure S2). *Pax3* at P4 and *Dbx1* at P14 are observed expressed in the superficial layers (Figure S2) suggesting that the IP8 population consists of superficial neuron progenitors.

**Figure 4.**
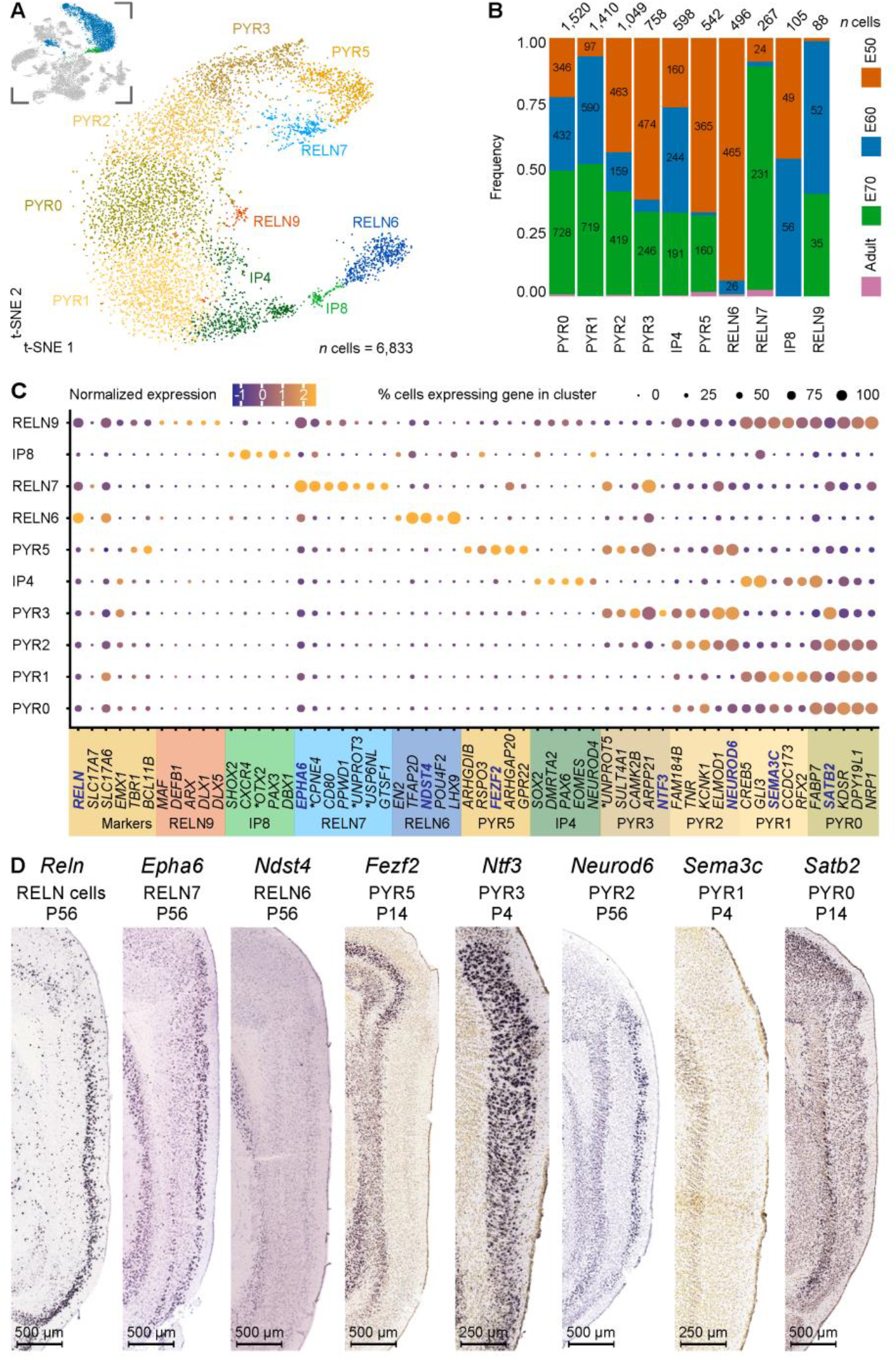
Subclustering of the excitatory neurons and intermediate progenitors of the developing entorhinal cortex (EC) highlight three *RELN*^+^ clusters and five pyramidal neuron clusters. **(A)** A t-SNE plot highlights subclustering of excitatory neurons and intermediate progenitors (number (n) of cells = 6,833) results in ten distinct clusters. The inserted image shows the excitatory neuron clusters highlighted in blue and intermediate progenitors in green, that were subclustered from the parent dataset. **(B)** Distribution of developmental stages within the ten clusters highlights five populations persist postnatally (three pyramidal neurons, PYR0, PYR2, PYR5 and two Reelin+ cell populations, RELN6, RELN7). Three IP populations were identified IP4, IP8 and RELN9 which the latter also expressed IN markers and RELN. IP4 was also detected postnatally but expressed IP makers *EOMES, PAX6* and *SOX2* **(C)** Unique gene signatures for the ten clusters were determined following differential gene expression analysis. Localization of genes with names in blue are shown in (D). * marks the human ortholog names (Table S1). **(D)** Sagittal sections from the ISH datasets from the Allen Developing Mouse Brain Atlas and Allen Mouse Brain Atlas highlight the location of selected markers from the RELN and PYR clusters across the EC layers at different developmental stages and in the adult (“Allen Developing Mouse Brain Atlas,” 2008; “Allen Mouse Brain Atlas,” 2004; Lein et al., 2007). Image credit: Allen Institute. Scale bars: 250 μm (for E18.5 and P4) and 500 μm (for P56).

An analysis of canonical genes for immature neurons with a pyramidal-like identity revealed five clusters. Clusters PYR0, PYR1, PYR2, PYR3 and PYR5 were tangentially located on the t-SNE plot (Figure 4A) and expressed low levels of *RELN* (Figure 4C). Surprisingly, we did not observe in our dataset any neuron populations expressing *CALB1*, a common pyramidal neuron marker reported in rats, mice and humans within the EC (Beall & Lewis, 1992; Diekmann, Nitsch, & Ohm, 1994; Fujimaru & Kosaka, 1996). *CALB1* may not be a suitable marker for pyramidal neurons in the pig EC as a previous study has reported the absence of expression in pyramidal neurons in the hippocampal region (Holm, Geneser, Zimmer, & Baimbridge, 1990). Clusters PYR0 and PYR1 shared similar transcriptional profiles but diverged in *NEUROD1* expression. Clusters PYR2 and PYR3 were also similar, but had divergent *ARPP21* expression. Only PYR1 and PYR3 could be detected in the prenatal brain and not in the postnatal brain. Given the similar profiles, it is likely that PYR0, PYR2 and PYR5 are adult populations. This does not explain however why all five populations are identified during early development, so we cannot eliminate the possibility that these are potentially different pyramidal neurons. Therefore, we deduce to have captured at least three pyramidal neuron populations. An analysis of mature pyramidal neuron markers only partially confirmed the notion of maturation of PYR1, PYR3 and PYR5, as *SLC17A6/7* (vGLUT1/2 genes) expression was observed in PYR0 and PYR2 but not PYR5 (Figure 4C). Validation of two unique gene markers in the Allen Developing Brain Mouse Atlas (“Allen Developing Mouse Brain Atlas,” 2008) corroborated the putative location of these cells. The gene *TNR* is highly expressed in PYR2 and PYR3 and was found in the superficial layers of the mouse EC (Figure S2). The gene *NEUROD6* was highly upregulated in PYR2, PYR3 and PYR5 and can be detected in both the superficial and deeper layers. Our investigations of the pig EC show BCL11B is expressed both in LII and LV of the EC (Liu et al., 2021)) and we found *BCL11B* to be highly upregulated in PYR5 suggesting that cluster PYR5 may reside in both the superficial and deep layers. Taken together, we have identified five pyramidal neuron subtypes during development and at least three mature pyramidal neuron types in the postnatal EC that are uniquely placed across the EC layers.

In addition to the pyramidal neurons, the EC harbors two types of principal neurons expressing Reelin. These are the stellate and fan cells that reside in LII/III in the MEC and LEC, respectively (Witter et al., 2017). Our subclustering analysis of the MEC revealed three distinct neuron populations that expressed *RELN* (RELN 6, RELN7 and RELN9), two of which were present postnatally (RELN6 and RELN7) (Figure 4 A,B). Apart from the known expression of Reelin and absence of CALB1 (at least in a proportion of Reelin neurons), few other genes are known to be expressed in these cells (Fuchs et al., 2016; Kitamura et al., 2014; Winterer et al., 2017). RELN9 was only present during gestation (mostly at E60-E70) (Figure 4B) and clustered closely to PYR0 and IP4. It expressed *DLX1, DLX5* and *ARX* (Figure 4A-C), together with *GAD1/2* (Figure S1B), suggesting it has more of a GABAergic IN progenitor (Friocourt & Parnavelas, 2011; Pla et al., 2018; Y. Wang et al., 2010)-like identity and we deduce this not to be a reelin excitatory neuron.

This left us with clusters RELN6 and RELN7, which lacked the pyramidal transcription factor, *EMX1(Chan* et al., 2001) (Figure 4C). These two clusters were expressed throughout all sampled time points and persisted in the postnatal adult brain (Figure 4B). They had distinct transcriptional profiles and were distinctly separated on the t-SNE plot (Figure 4A, B). Interestingly, RELN6 cells are more abundant than RELN7 cells at E50 suggesting that more of this RELN subtype is born during early neurogenesis (Figure 4B). In contrast, more RELN7 cells were captured at E70 (late neurogenesis in the pig (Liu et al., 2021)) than RELN6 cells (Figure 4B). When assessing unique markers from these clusters in the Allen mouse atlases, the RELN6 uniquely identified gene, *Lhx9*, was expressed across all the EC layers but also across the parahippocampal cortex at E18.5 and P14 in the mouse (“Allen Developing Mouse Brain Atlas,” 2008) (Figure S2). This marker is known to be expressed in Reelin negative (-) pioneer neurons across the cortex (Bertuzzi et al., 1999) and is therefore not an ideal marker for identifying the RELN6 SC within the MEC alone. In the Allen Developing Mouse Brain Atlas, another marker, *Ndst4* was expressed at higher levels in the superficial layers at P56 in the mouse EC (“Allen Developing Mouse Brain Atlas,” 2008) (Figure 4D) which indicates their location within the superficial layers. When assessing genes from cluster RELN7 in the Allen Developing Mouse Brain Atlas, moderate levels of *Reln* and *Cpne4*, which are both expressed in the superficial layers of the mouse EC and were more enriched in the LEC than the MEC LII (“Allen Developing Mouse Brain Atlas,” 2008) (Figure 4C, D). Further, *Epha6* was upregulated in cluster RELN7 and could be observed in the LII-LIII in the Allen Mouse Brain Atlas (“Allen Mouse Brain Atlas,” 2004), confirming the RELN7 neurons are also located in the superficial layer. To sum, two *RELN* excitatory neuron populations were identified which persisted after birth.

### Induced expression of six transcription factors in human iPSCs results in stellate cell-like progenitors

The Reelin+ SCs are a vulnerable cell population in the EC (Kobro-Flatmoen et al., 2016; Leng et al., 2021) and would be an interesting cell type to model in-vitro for studying AD. We sought to use our scRNA-seq data to produce a novel protocol for the production of SCs from pluripotent stem cells by induced overexpression of SC specific transcription factors. Therefore, looked for candidate TFs among the twenty most enriched TFs in the two *RELN* populations based on the TFcheckpoint list of DNA-binding RNA polymerase II TFs curated by Chawla *et. al* (Chawla, Tripathi, Thommesen, Laegreid, & Kuiper, 2013) using the Findmarkers tool in Seurat (Figure 5A). We focused our attention on the RELN7 cluster, given it represented a mature SC profile that was located in the superficial layers of the EC. *RORB* was found to be one of the twenty differentially expressed TFs, and it has recently been identified as a marker for vulnerable EC excitatory neurons (Leng et al., 2021). Of the 20 TFs we focused on six, *RUNX1A1, SOX5, FOXP1, MEF2C, TCF4* and *EYA2* based on a literature search showing prominent roles for these TFs in neurodevelopment and differentiation. Runx1t1 is specifically expressed in neurons and plays a role in hippocampal neuron differentiation (Linqing et al., 2015; Zou, Li, Han, Qin, & Song, 2020), which also lies in close proximity with the EC within the ventral telencephalon and both structures arise from the medial pallium (Abellan, Desfilis, & Medina, 2014). Eya2 is expressed in neural progenitors and is repressed by Foxg1 during corticogenesis, playing an important role in neuronal differentiation (Kumamoto et al., 2013). Foxp1 is co-expressed with Satb2 and detected in LIII-LVa neurons (Hisaoka, Nakamura, Senba, & Morikawa, 2010). It is also implicated in neural stem cell differentiation and neuronal morphogenesis (Braccioli et al., 2017; X. Li et al., 2015). We recently demonstrated that MEC SC express SATB2 which further raised our interest in FOXP1 (Liu et al., 2021). *SATB2* was also detected in both RELN7 and RELN9 (Figure 4C). SOX5 is an important regulator of early-born deep layer neurons (Kwan et al., 2008) and in differentiation of specific corticofugal neuron subtypes (Lai et al., 2008) but may be involved in SC fate given they also express the deep layer marker, BCL11B (Liu et al., 2021). *BCL11B* was also upregulated in the RELN7 population and more so than RELN6 (Figure 4C). Tcf4 is expressed in cortical and hippocampal neurons, highly enriched in the pallial region (Kim, Berens, Ochandarena, & Philpot, 2020) and is involved in neurogenesis, neuronal differentiation and hippocampal formation (Hill et al., 2017; Mesman, Bakker, & Smidt, 2020; Schoof et al., 2020). It is also implicated in memory and spatial learning (Kennedy et al., 2016). Mef2c plays a role in neuronal development and is expressed in hippocampal neurons (Adachi, Lin, Pranav, & Monteggia, 2016). It is also implicated in cognitive learning, including working memory and object recognition (Mitchell et al., 2018). An assessment of these genes in the mouse brain ISH data from the Allen Mouse Brain Atlas demonstrated that both *Tcf4* and*Mef2c* are expressed in the superficial layers of the EC, whereas the remaining genes were expressed across all the layers (“Allen Mouse Brain Atlas,” 2004) (Figure 5B). The sequences of these six TFs were also available from Addgene and cloned into plasmids from either mouse (Foxp1, Sox5) or human (TCF4, MEF2C, RUNX1T1, EYA2) RNA.

**Figure 5.**
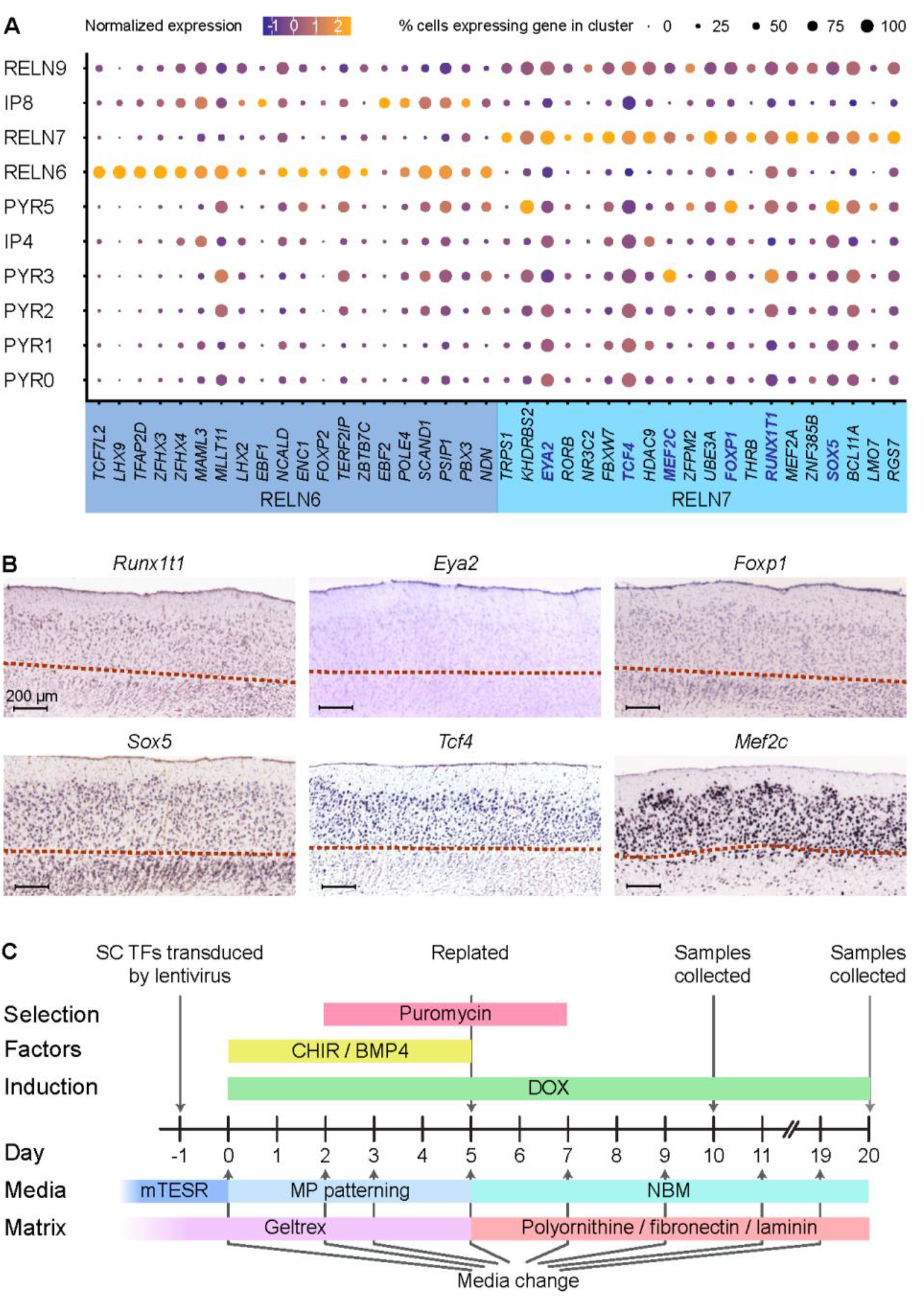
Unique transcriptional factors upregulated in *RELN* positive clusters. **(A)** Expression of the top twenty enriched transcription factors (TFs, x-axis) were identified for each of the *RELN-positive* and putative stellate cell (SC) populations (RELN6, 7) and are visualized across the identified progenitor and excitatory neuron populations (y-axis) in the developing EC. The selected genes for reprogramming induced pluripotent stem cells (iPSCs) are highlighted in blue within the RELN7 cluster. **(B)** The laminar expression of the six SC highlighted TFs (*Runx1t1, Eya2, Foxp1, Sox5, Tcf4*, and *Mef2c*) was investigated by ISH in sagittal sections of the publically available Allen Mouse Brain Atlas (“Allen Mouse Brain Atlas,” 2004; Lein et al., 2007). Image credit: Allen Institute. The red dotted lines demarcate the *lamina dissecans*. **(C)** Schematic diagram of the protocol used for directly reprogramming iPSCs into SCs using the selected TFs, including timing of media and matrix change, transfection, gene induction (using DOX), addition of patterning factors (CHIR 99021 and BMP4), and collection of samples. Abbreviations: Doxycycline (DOX); pluripotent stem cell media (mTESR); neurobasal medium with medial pallium patterning factors (MP patterning); neurobasal medium for maintenance of neurons (NBM).

In order to produce SC from human iPSCs, we applied an overexpression approach in combination with culture conditions that induced a medial pallial fate. Evidence suggests the MEC is derived from the medial pallium (Abellan et al., 2014; Bruce & Neary, 1995; Desfilis, Abellan, Sentandreu, & Medina, 2018). We also tested whether Ngn2 was necessary in the direct programming approach since Chen *et al*. have demonstrated that patterning neurons with specific factors in combination with ectopic Ngn2 expression yielded a more uniform population with a specific CNS fate (Chen et al., 2020). We performed lentiviral transduction in the human iPSC lines, SFC 180-01-01 (SFC) and WTSli024A (WTS) using different combinations of lentiviruses expressing the TFs in the presence or absence of Ngn2 (applying a leave-one-out approach in the presence of Ngn2) and cultured the transduced cells in CHIR and BMP4 for 5 days (Figure 5C). Puromycin selection was initiated on day (D)2 and continued 5 days to eliminate non-transduced cells. The cells were re-seeded on D5 and propagated in neuronal medium for up to a further 15 days (total 20 days) (Figure 5C). We also included a negative control where the human iPSCs were not transduced with TFs. We determined that these cells were unable to differentiate into a neuronal fate in the medial pallial culture conditions alone (Figure 6). As to be expected, the Ngn2 alone treatment resulted in neuronal morphology at D10 and D20, as has been described previously (Y. Zhang et al., 2013). The combination of the six SC TFs in the presence of Ngn2 also resulted in neurons with axonal projections at D20 (Figure 6A). However, this was only observed in the SFC line, and pronounced cell death was observed in the WTS cell line, which might be attributed to cell line-specific toxicity. Strikingly, the combination of the six SC TFs in the absence of Ngn2 resulted in bipolar cells at D10, and at D20 a more neuron-like morphology was observed (Figure 6A). The leave-one-out SC TF approach in the presence of Ngn2 also resulted in neurons with extending dendrites at D20 (Figure S3A). By morphological observation alone, we were not able to exclude the contribution of any particular, or combinations of SC TF in directing a neuronal fate in the presence of Ngn2.

**Figure 6.**
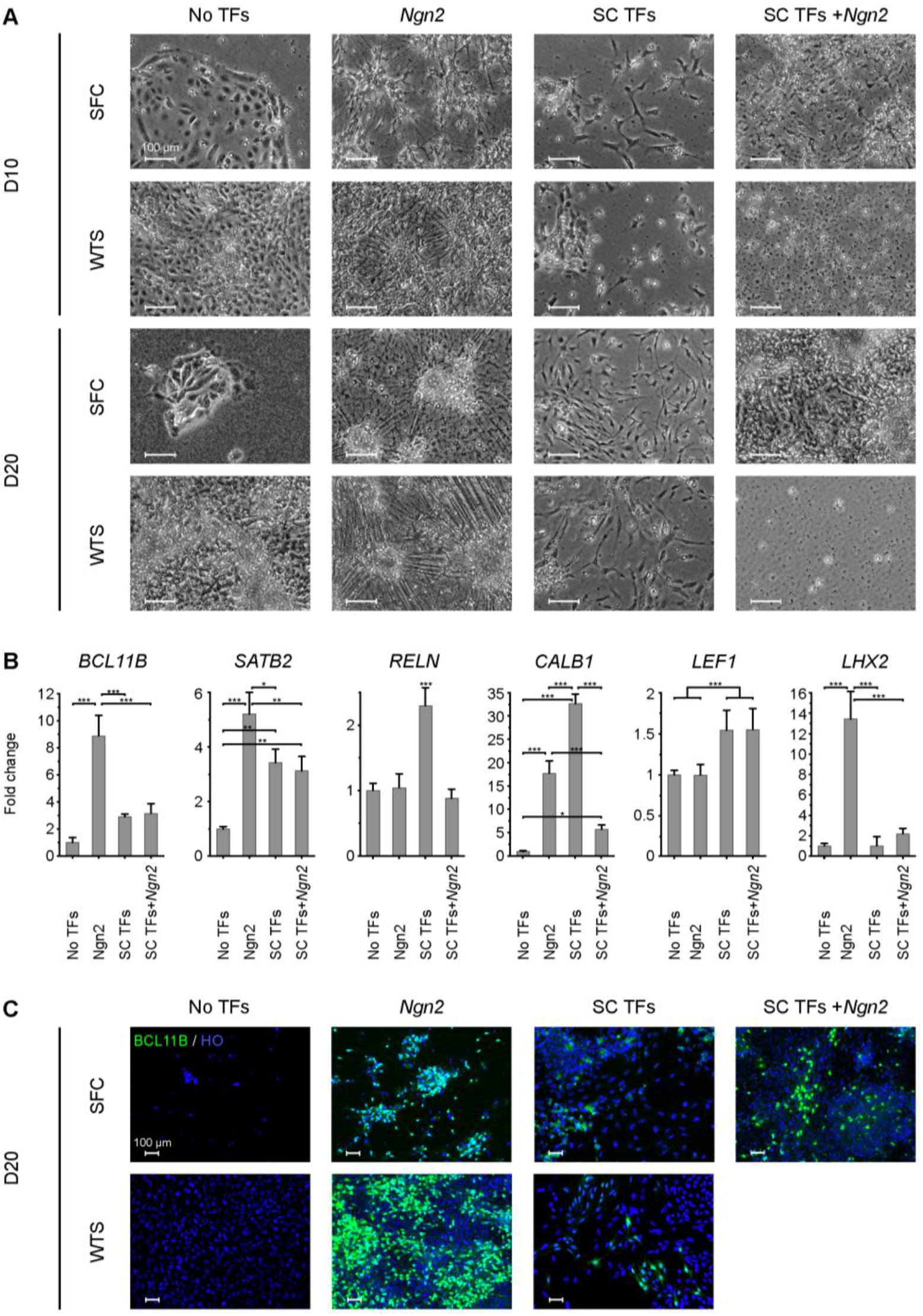
Direct reprogramming of induced pluripotent stem cells (iPSCs) results in the production of stellate cell-like cells. **(A)** The morphological changes of the two iPSC lines SFC 180-01-01 (SFC) and WTSli024A (WTS) are shown after transduction with the stellate cell transcription factors alone (SC TFs), the SC TFs plus *Ngn2, Ngn2* alone and without any TFs at Day (D)10 and D20 post transduction. Scale bar: 100 μm. (B) RT-qPCR analysis of pooled D10 and D20 samples highlights upregulated expression of *RELN, SATB2, CALB1* and *LEF1* in SC TF transduced iPSCs compared with iPSCs not transduced with any TFs. Expression values were normalized to *GAPDH* and presented as fold-change relative to No TFs. Bars represent the mean values of three replicates. The error bars represent the SD. Statistical one-way ANOVA, with Tukey’s multiple comparisons test was performed and significance is reported as: *p < 0.05, **p < 0.01, ***p <0.001. **(C)** Expression of BCL11B was determined in the transduced SFC and WTS iPSCs, including the SC TFs alone treatment, at D20. HO; Hoechst. Scale bar: 100 μm.

In order to further elucidate the identity of the neurons produced, we investigated the expression of known SC markers, *BCL11B, SATB2, RELN*, and genes upregulated in the MEC LII, *LEF1* and *LHX2* (Abellan et al., 2014; Hebert, Mishina, & McConnell, 2002; Liu et al., 2021; Medina, Abellan, & Desfilis, 2017) in the cells reprogrammed with all six SC TFs with or without Ngn2, compared to Ngn2 alone, at D10 and D20. We selected *BCL11B* since we have determined earlier that most earlyborn SC are BCL11B+ (Liu et al., 2021), however other SC populations are BCL11B-. The qPCR confirmed expression of *BCL11B* in all three treatments, but the highest levels of expression were observed in the Ngn2-only induced cells (Figure 6B). A significant increase in *SATB2* expression was also observed in all three treatments, but was highest in the Ngn2-only neurons (Figure 6B). Interestingly, a significant increase (p <0.001) in *RELN* expression was observed only in the cells induced with expression of the SC TFs alone (Figure 6B). Together, the SC TFs induced co-expression of *BCL11B, SATB2*, and *RELN*, mirroring the unique molecular profile of EC SCs. To rule out a pyramidal neuron identity we analyzed the cells for *CALB1*. Of interest, was a high expression of the pyramidal marker, *CALB1* in the cells induced with the SC TFs alone. This was unexpected, as only a smaller subpopulation of entorhinal SCs, the intermediate SCs, express both Reelin and Calbindin (Witter et al., 2017). Further, *LEF1* was upregulated in both treatments containing SC TFs but not in the Ngn2 alone treatment (Figure 6B). In contrast, *LHX2* was only upregulated in the Ngn2-only treatment. The analyses indicate the SC TFs were able to induce the expression of *BCL11B, SATB2, RELN, CALB1* and *LEF1* in the human iPSCs derived neurons by D20. To further determine a SC glutamatergic identity, we assessed the iPSC-derived cell populations at D10 and D20 for expression of BCL11B, Reelin, and vGLUT1 by performing immunocytochemistry. Interestingly, none of the neurons from the different treatments expressed Reelin or vGLUT1 (data not shown). The absence of Reelin was not surprising since expression of this gene is switched on later in development. However, BCL11B expression was observed in cells transduced with Ngn2, SC TFs and Ngn2 and SC TFs alone, at D20 (Figure 6C) and as early as D10 in the WTS line (Figure S3B). This observation indicates that the SC TFs alone were able to induce BCL11B expression in cells with neuronal morphology. Omission of one of the six SC TFs in the presence of Ngn2 also resulted in BCL11B expression in the cells (Figure S4). To sum, the SC TFs alone were able to differentiate human iPSCs into SC-like cells with neuronal processes and several unique SC markers. Further, *RELN* was only detectable at the RNA level which suggests an immature neuronal phenotype given the lack of vGLUT1 expression.

## Discussion

Our study demonstrates the cellular architecture of the developing EC and provides a proof-of-concept approach for using scRNA-seq data to produce region-specific neurons from induced pluripotent stem cells. We demonstrate that scRNA-seq data from the developing and postnatal brain can be used to identify transcriptional regulators important for specification of neuronal cell subtypes. We identified in a proof-of-concept study that six TFs: *FOXP1, SOX5, TCF4, MEF2C, RUNX1T1* and *EYA2*, when added in combination, resulted in the differentiation of entorhinal SC-like cells. Further studies are required to identify if one or more of these TFs may be redundant in the reprogramming process. This could have been facilitated in hindsight if Ngn2 had been removed from the leave-one-out experiment. Nevertheless, we prove that transcriptional regulators of neuronal subtype cell fate can be identified from scRNA-seq clusters which are defined in the curated TFcheckpoint list and extracted using the Findmarkers tool in Seurat. This study will fuel future studies for streamlining the reprogramming process and optimizing the transduction process.

The phenotype of the SC-like cells produced may reflect the intermediate SC which is an entorhinal cell type that expresses both Reelin and Calb (Fuchs et al., 2016; Witter et al., 2017). Interestingly, we could not detect *CALB1* in the RELN7 cluster, which could be attributed to low gene expression and threshold cut-offs of low expressing genes. Our own and other studies have shown peculiarities of CALB expression in the pig ventral telencephalon. For example, CALB1 protein is localized only in white matter tracts in the EC of the postnatal pig EC and could not be detected at all, prior to E100 (two weeks prior to birth) (Liu et al., 2021). Further inherent differences show a lack of CALB1 in mossy fibers, granule cells and pyramidal neurons in the pig hippocampus (Holm et al., 1990). Post transcriptional mechanisms might be responsible for a lack of protein expression in pyramidal neurons in the EC, such as upstream repressors of the open reading frame (Calvo, Pagliarini, & Mootha, 2009), frameshifting events (Dinman, 2012) or alternative transcription start sites that create different transcript isoforms (X. Wang, Hou, Quedenau, & Chen, 2016). The upregulation of *RELN* expression in the SC-like cells points to a SC phenotype as it is rarely expressed in neurons. An exception to the entorhinal SC are the glutamatergic granule cells in the cerebellum (Pesold et al., 1998). Instead *RELN* is predominantly expressed in GABAergic interneurons (Pesold et al., 1998), but the combination of genes in these cells did not reflect an interneuron phenotype. Instead, the coexpression of BCL11B and SATB2 highly favors a superficial or deep layer entorhinal neuron phenotype in the MEC (Liu et al., 2021), however LEF1 is highly enriched in the MEC LII (Ramsden, Surmeli, McDonagh, & Nolan, 2015) which points the cells towards having a superficial layer identity. The lack of Reelin protein expression is not surprising since we see protein expression of Reelin turns on several days (approx. 10 days in the pig) after birth of the SC (Liu et al., 2021), which may even be longer in the human, given the longer gestation period. Prolonged cultures of these iPSC-derived neurons and more extensive marker evaluation could help to confirm their identity. We cannot rule out that the RELN6 population are also stellate cells residing in the superficial layers. These cells also expressed *BCL11B* but were *RORB* negative and expressed only low levels of *SATB2*. Our previous studies in the mouse EC show Reelin+/Bcl11b+ neurons are born in the superficial layers (LII and LIII) from E12.5 to E15, but most were born at E12.5 (Liu et al., 2021). Another subset of Reln+/Bcl11b-were also born from E12.5 to E15 (Liu et al., 2021). In the pig scRNA-seq data, both RELN6 and RELN7 expressed similar low levels of *BCL11B* but appear to be born at different time points when assessing the varying proportions of the cells captured at E50 and E70. This suggests RELN6 cells may arise earlier than RELN7 cells. However, we did not capture a *RELN*+ population expressing high levels of *BCL11B* in the postnatal brain samples so it is difficult to confer our previous mouse data to our current porcine scRNA-seq data. Further studies analyzing differences between the RELN populations may help to distinguish the RELN subtypes better.

Our study also demonstrates that pig scRNA-seq data can be readily applied for generation of novel protocols in human iPSCs. Indeed we found the developing EC dataset projected extremely well using scmap to human and mouse fetal datasets. This was the case, even when the human datasets were at unmatched gestational ages and from the embryonic prefrontal cortex and adult middle temporal gyrus. Our previous findings have also identified that the pig poses as an excellent model of neurogenesis, given its long gestational length, completion of corticogenesis prior to birth and presence of a moderate outer subventricular zone (Liu et al., 2021).

Our single-cell profiling of the porcine postnatal EC revealed a high quality dataset which corroborates and further enhances knowledge of EC cell types in the postnatal brain. Another scRNA-seq study was recently published on the human adult EC from both AD patient and healthy brains, however, their data showed a median of 646 genes captured in 13,214 sequenced cells (Grubman et al., 2019) Similarly, Leng et al., profiled a higher number of cells in the human caudal EC (42,528 cells) however the depth of sequencing resulted in a variable number of genes captured per cell, for e.g. a mean of 2,000 genes per excitatory cell) (Leng et al., 2021). Our dataset resulted in a mean of 2,798 different genes captured per cell in the 24,294 cells. In a report by Grubman et al., the vast majority of cells captured in their dataset were predominantly glia with very few neurons (Grubman et al., 2019). For example in their study, only six neuronal subtypes could be identified in the adult EC. The Leng et al. study extended this knowledge and identified nine excitatory neuron populations. Our study identified twelve excitatory neurons, some of which exist only during development, but furthers the field’s knowledge on the EC subtypes. We also characterized the neurons based on the two main neuron types of the EC, the pyramidal and SC subtypes. We identified at least three subtypes of pyramidal neurons and two stellate neuron subtypes in the postnatal MEC. We also identified four IN subtypes in the postnatal MEC which is fewer than the nine inhibitory neuron populations identified in Leng et al., in the adult human brain (Leng et al., 2021) but were able to detect six IN progenitor populations, when also including the *RELN*+ population that clustered closely to the excitatory neurons in the excitatory neuron subclustering analyses. The variation in numbers of IN populations seen in our study compared to the Leng study may be due to species-specific differences, sampling differences or differences in clustering methodology. In regions where IN diversity have been well studied such as the amygdala, at least 6 classes of INs have been identified based on their discharge properties and electrophysiological profiles in the basolateral amygdala (Polepalli, Gooch, & Sah, 2020) and similar research is required in the EC to fully differentiate between the different IN subclasses. The INs lying in the superficial layers are particularly interesting for researchers studying the mechanisms underlying grid cell firing. We found expression of unique genes from IN4 and IN0 to be highly abundant in the superficial layers of the mouse EC, but of the two populations, only IN4 is SST-which suggests this may be the IN population of most interest, since SST-/PVALB+ INs are directly coupled to grid cell firing (Miao, Cao, Moser, & Moser, 2017). We also confer with the Leng et al. study, that two oligodendrocyte populations exist in the postnatal brain (Leng et al., 2021). The two previous scRNA-seq studies on the EC have concentrated on differences between healthy and diseased brains and characterization of vulnerable cell populations to AD rather than on the characterization of the cell types *per se*. Our dataset therefore provides excellent insight into the cell diversity of the EC due to the increased cell capture rate, depth of sequencing, and subclustering analyses.

In conclusion, we demonstrate that scRNA-seq data from the developing brain can be used to produce novel protocols for direct programming of pluripotent stem cells into subtype specific entorhinal cell fates and we provide evidence in a proof-of-concept approach to produce SC-like cells. Production of novel cell types ex-vivo will facilitate research into tissues, such as the EC, which is afflicted early on in e.g. AD. We envisage future studies could further build upon this dataset by adding the spatial dimension and neuron connectivity to build even more accurate and detailed maps of the spatial processing system.

## METHODS

### Aim, design and setting of the study

#### Animal welfare and collection of brains

The experiments conform to the relevant regulatory standards for use of fetal material. Crossbred male and female pig fetuses at Embryonic day (E)50, E60, and E70 of development were obtained from inseminated Danish Landrace/Yorkshire sows with Duroc boar semen from a local pig production farm. Deceased postnatal pigs were obtained at postnatal day (P)75 as a gift from Per Torp Sangild at the University of Copenhagen. Adult brains were obtained from sows killed for another study using an overdose of sodium phenobarbital by a professional issued with a license from the Danish Animal Experiment Inspectorate.

#### Single-cell preparation

In total we prepared 10 sequencing-libraries form isolated MECs from E50 (whole cell, 3 brains, 1 batch), E60 (whole cell, 4 brains, 3 batches), E70 (whole cell, 4 brains, 3 batches; nucleus 1 brain, one batch) and from adult sow MEC (whole cell, 1 brain, 1 batch; nucleus, 1 brain, 1 batch) (Table S1). Briefly, the individual MEC tissue was digested using a papain dissociation method, according to the manufacturer’s guidelines (Worthington) with small modifications.

The EC was macroscopically dissected out (approx. 1 mm^3^) in the digest medium (1x PBS (Thermo Fisher Scientific), 1x Penicillin-Streptomycin (Sigma-Aldrich)), transferred to a 3.5cm Petri dish and incubated in 1mL papain solution for 30 min at 37°C. The tissue was gently triturated 20 times. The cell suspension was diluted with 1 mL FBS (BioWest) and centrifuged for 5 min at 300 g at room temperature (RT). The supernatant was discarded and the cell pellet was resuspended in 2.7 mL digestion media (1x Neurobasal medium (Thermo Fisher Scientific), 10% FBS (BioWest), 1x Penicillin-Streptomycin (Sigma-Aldrich)), 300 uL albumin-ovomucoid inhibitor, and 150 uL DNAse solution. The cell suspension was carefully layered on top of 5 ml of albumin-inhibitor solution in a 15 mL falcon tube and centrifuged for 6 min at 70 g at RT. The supernatant was discarded and the cell pellet was resuspended in 5 mL cell-resuspension media (1x Neurobasal medium (Thermo Fisher Scientific), B27 (Thermo Fisher Scientific), 1x Penicillin-Streptomycin (Sigma-Aldrich), bFGF (5 ng/ml, Prospec). The cells were counted (NucleoCounter, ChemoMetec) and ranged in viability from 80.4 - 88.4% (Viability and Cell Count assay), diluted to 100-2000 cells/uL used for single-cell library preparation.

#### Single-nuclei isolation

Nuclei extraction was performed as described before (Krishnaswami et al., 2016) with the following modifications (Pfisterer et al., 2020). Prior to nuclei extraction, nuclei isolation medium 1 (NIM1) (250 mM sucrose, 25 mM KCl, 5 mM MgCl2, 10 mM Tris Buffer pH8), NIM2 (NIM1 buffer supplemented with 1 μM DTT (Thermo Fisher Scientific) and 1x EDTA-free protease inhibitors (Roche) and homogenization (NIM2 buffer supplemented with Recombinant RNase Inhibitor (0.4 U/μL, Takara), SUPERase in (0.2 U/μL, Thermo Fisher Scientific) and Triton (0.1% v/v, Sigma-Aldrich)) buffers were prepared. Briefly, sectioned frozen brain tissue was placed into pre-cooled 1ml dounce homogenizer (Wheaton) with 1ml ice-cooled homogenization buffer. Tissue was dissociated on ice using 5-6 strokes with the loose pestle and 15-17 strokes with the tight pestle. Homogenate was first filtered through a 70 μm filter. Nuclei were collected (900 g, 10 min) and resuspended in 500μl staining buffer (nuclease free PBS (1X, Thermo Fisher Scientific), BSA (0.5% wt/vol, Sigma-Aldrich), SUPERase in (0.2 U/μL, Thermo Fisher Scientific)). The sample was stained with 7-AAD (2ug/uL, Sigma-Aldrich) in order to visualize nuclei during FACS sorting. 7-AAD positive nuclei were FACS-isolated (70 μm nozzle, BD Biosciences, BD FACSAria™) into a 1.5ml eppendorf tube containing 10 μl 10% nuclease free BSA (Thermo Fisher Scientific). Single 7-AAD+ nuclei were isolated using the gating strategy similar to Pfisterer et al (Pfisterer et al., 2020).

### Single-nuclei RNA-seq library preparation and sequencing

Whole cells were loaded onto the 10X Genomics microfluidic chip according to the Chromium Single Cell 3’ Reagent Kits User Guide version 2 chemistry (10X Genomics). The single-nuclei samples were loaded as per whole cells, albeit the samples were not diluted. 10-12.000 thousand cells/nuclei were loaded from each sample.

Libraries from two samples at different stages were pooled and sequenced together on an Illumina NextSeq 500 (Table S1) following the NextSeq System Denature and Dilute Libraries Guide Protocol A: Standard Normalization Method (illumina). The NextSeq 500/550 High Output Reagent Cartridge v2 75 cycles (illumina) kit was used for the whole cell and single-nuclei samples and the pooled library were sequenced on NextSeq 500. The sequencing cycles were: Read1, 26 cycles, i7 Index 8 cycles, i5 Index 0 cycles; Read2, 57 cycles.

### Plasmid design and construction

The selected genes of interest (GOIs) were cloned into the pTet-O-Ngn2-puro plasmid (Addgene #52047) by replacing *Ngn2* with the GOI. This ensured that the GOI was fused to the puromycin resistance gene. Plasmids containing *EYA2* (Addgene #49264), RUNX1T1 (Addgene #49264), *TCF4* (Addgene #109144), *Foxp1* (Addgene #16362), *Sox5* (Addgene #48707) and *MEF2C* (Addgene #61538) were all acquired from Addgene. Forward and reverse primers were designed for subcloning all GOIs (with the exception of MEF2C which was already subcloned into a lentiviral Tet-O plasmid (Addgene #61538) (see Table S2 for primer sequences). Gene inserts were amplified from the plasmids by PCR amplification using 400 - 600 ng template plasmid, 0.5 μM forward and reverse primer (Eurofins Genomics), 5 μL Pfu DNA Polymerase 10X Buffer (Promega), 1 μL Pfu DNA Polymerase (Promega), and 200 μM dNTP mix (Promega) in a 50 μL reaction volume. The following program was used in a PTC-200 Thermal Cycler (MJ Research): 1 cycle of 2 min at 95 °C, 40 cycles of 1 min at 95 °C, 30 sec at 60 °C, and 1 min at 72 °C. The PCR amplified inserts were run on a 1% agarose gel and purified using the Wizard SV Gel and PCR Clean-Up System kit (Promega) according to the manufacturer’s instructions. The gene insert was then ligated into a Zero Blunt TOPO vector using the Zero Blunt TOPO PCR Cloning Kit (Invitrogen) according to the manufacturer’s protocol. The TOPO vector containing the gene insert was heat-shock transformed into competent E. coli (NEB Stable C3040I), and grown at 37 °C O/N on agar (Sigma-Aldrich) plates containing 50 mg/mL kanamycin (Sigma-Aldrich) for selection. Individual colonies were propagated at 37 °C O/N in 5 mL Luria-Bertani (LB) broth (Sigma-Aldrich) supplemented with 50 mg/mL kanamycin. Plasmids were isolated using the PureYield Plasmid Miniprep System (Promega) according to the manufacturer’s protocol. The GOIs were isolated from the purified plasmids by digestion with EcoRI (Esp3I for RUNX1T1) and Xbal according to the manufacturer’s protocol (FastDigest, Thermo Fisher Scientific). Positive clones were purified from a 1% agarose gel. The GOI was then inserted and ligated into a dephosphorylated pTet-O-Ngn2-puro (addgene 52047) backbone to create the final plasmid; pTet-O-GOI-puro using the LigaFast Rapid DNA Ligation System (Promega) and heat-shock transformed into competent E. coli. Individual colonies were selected and isolated using the PureYield Plasmid Miniprep System (Promega) according to the manufacturer’s protocol. Correct plasmid inserts were validated by digestion with EcoRI and XbaI (FastDigest, Thermo Scientific). The plasmids were verified by Sanger sequencing (Eurofins Genomics) and stored at -80 °C until use for virus production.

#### Lentiviral production and titering

Second generation lentiviruses were produced following transfection of HEK293FT cells. On the day of transfection, HEK293FT cells were cultured in fresh DMEM supplemented with 10% newborn bovine calf serum (NCS) (Hyclone, New Zealand) in the absence of antibiotics. The cells were transfected as follows: In one tube, 50 μL Lipofectamine 3000 reagent (Invitrogen) was diluted in 1.5 mL Opti-MEM (Gibco; Thermo Fisher Scientific). 40 μL P3000 reagent (Invitrogen) and 5 pmol DNA plasmids (1.5 pmol psPAX2 (Addgene #12260), 1.5 pmol pMD2.G (Addgene #12259) and 2 pmol transfer plasmid containing the GOI) were diluted in 1.5 mL Opti-MEM. The two tubes were mixed and incubated at RT for 10 min and the solution was then added dropwise to the cells. After 24 h the media was replaced with fresh DMEM supplemented with 10% NCS and incubated for 48 h. The virus was harvested 72 h post transfection. The virus was spun at 500 g for 5 min and the supernatant filtered through a 45 μm polyethersulfone filter (Fisher Scientific) and subsequently concentrated, by spinning at 23,000 rpm (using rotor Beckman Type 60 Ti) for 2 h at 4 °C. The supernatant was discarded and 100 μL PBS was added to the pellet and incubated for 1 h at 4 °C. The pellet was resuspended by pipetting and stored at -80 °C until use for transduction.

Titering of virus was performed on HEK293FT cells by adding lentivirus at different amounts (from 1 to 15 μL virus). A virus with a predetermined titer was added as a reference and a negative control without virus was included. On Day 1, the media was substituted with fresh media, and 48 h (Day 2) after transduction, the genomic DNA (gDNA) was extracted from the cells using the PureLink Genomic DNA Mini Kit (Invitrogen) according to the manufacturer’s protocol. Each sample was analyzed in triplicate by qPCR using both primers for a sequence fused to the GOI (LV2), fused to all GOIs, and for the β-actin intron (see Table S3 for primer sequences). The LV2 was used to determine the degree of viral integration into the genome. The β-actin gene was used to determine the number of genome copies, as an estimate for the total number of cells. For the qPCR reaction, 5 μL SYBR green (Roche), 1 μL 10 mM of both forward and reverse primer and 15 ng gDNA was mixed in a 10 μL reaction volume and loaded in a white qPCR plate with optical caps (Thermo Fisher Scientific). Samples were spun briefly and run on a Mx3005P qPCR machine (Agilent) using the following program: 1 cycle of 10 min at 95 °C, 40 cycles of 20 sec at 95 °C, 20 sec at 55 °C, and 30 sec at 72 °C, followed by a melting curve analysis.

The viral titers were determined based on a recently published approach with minor variations (Gill & Denham, 2020). Briefly Ct values were converted from their intrinsic exponential relation to linear related Xo values, using the Xo method (Thomsen, Solvsten, Linnet, Blechingberg, & Nielsen, 2010). The triplicate LV2 X o mean values for a sample was normalized to its mean Xo value for β-actin to obtain the copy number of integrated GOI relative to β-actin gene copies in the given sample. A standard curve of normalized LV2 sequence copy number versus the volume of virus that was added to the sample initially was fitted for each titration of the virus, respectively. All curves followed a linear relationship with R2 ranging from 0.889 to 0.995. The relative GOI integration for the reference virus was then calculated and inserted into the linear regression curves for each respective virus, where (VPD, volume of produced virus): VPD = a * relative copy number of reference virus + b. The VPD is a measure of the volume of the produced virus that is needed, under the same conditions as the reference virus, to obtain the same degree of GOI integration as was obtained for the reference virus. The VPD for each of the produced viruses was hence calculated by inserting the relative copy number of the reference virus in the linear regression for the respective virus produced. The titer of the produced virus was finally calculated based on the titer of the reference virus, using a multiplicity of infection (MOI) of 10, where U is units of viral particles: VPD = (Number of cells * MOI) / Titer (U / μL).

#### Cell culture

For direct reprogramming, two human iPSC lines were selected including the human SFC180-01-01 (female donor, healthy, age unknown) and WTSli024A (female donor, healthy, age 50-54) (both acquired from the EBiSC cell bank). The cells were maintained in Geltrex (Gibco; Thermo Fisher Scientific) coated dishes with mTESR media (Stemcell Technologies) supplemented with 1% Penicillin-Streptomycin (Gibco; Thermo Fisher Scientific). The cells were passaged using 0.5 mM EDTA (Merck) and propagated until they reached 50-60% confluency, before initiating reprogramming using the produced lentivirus. The cell lines were cultured at 37 °C in 5% CO2 with 90% humidity.

#### Transduction of human iPSCs

Human iPSCs were transduced in mTESR in the absence of antibiotics and with 6 μg/mL polybrene (Millipore). The lentiviruses containing the GOIs were transduced using 10 MOI in the presence of the reverse tetracycline transactivator M2rtTA (MOI 10). After 3 h the media was supplemented with half the usual volume of fresh media. The day after viral transduction was considered Day 0, and the media was exchanged for medial pallium (MP) patterning media (47% Neurobasal A medium, 47% DMEM/F12+Glutamax, 1% Insulin-Transferrin-Selenium A (Gibco; Thermo Fisher Scientific), 2% B-27 Supplement (Gibco; Thermo Fisher Scientific), 1% CTS N-2 supplement (Gibco; Thermo Fisher Scientific), 0.5% GlutaMAX (Thermo Fisher Scientific), 1% Pen Strep, 0.3% Glucose, 3 μM CHIR 99021 (Sigma-Aldrich), 5 nM BMP4 (R&D systems), 1 μg/mL doxycycline (Sigma-Aldrich)). On Day 2 and 3 the media was exchanged for MP patterning media supplemented with 3 μg/mL puromycin to select for transduced cells. On Day 5, the cells were dissociated using dispase (Sigma-Aldrich), spun for 5 min at 300 g and cultured in NBM media (96% Neurobasal medium, 2% B-27 Supplement, 1% GlutaMAX, 1% pen strep, 1 μg/mL doxycycline, 200 μM Ascorbic Acid (Sigma-Aldrich), 1 μM DAPT (Sigma-Aldrich)) and 3 μg/mL puromycin. Upon confluency, the cells were split onto triple coated (1x polyornithine (Sigma) 10 μg/mL fibronectin (Sigma-Aldrich),10 μg/mL PolyD laminin (Sigma-Aldrich) plates. The cells were propagated in the NBM medium up until D20.

#### Immunocytochemistry

The differentiated human iPSCs were fixed on glass coverslips (VWR, Denmark) in 4% PFA for 15 min at RT and stored at 4°C until use. The cells were washed in PBS and permeabilized in 0.25% Triton X-100 in PBS, washed once in PBS-T (0.1% Tween-20 (Sigma-Aldrich) in PBS) and then treated with blocking buffer (10% NDS, 3% BSA, 0.1% Tween-20 in PBS) for 30 min at RT. Immunocytochemistry was performed using a combination of primary antibodies targeted against BCL11B (1:400, Abcam, ab18465), Reelin (1:100, Santa-cruz, sc-25346) and vGLUT1 (1:200, Abcam, ab79774) diluted in blocking buffer 2 for 60 min at RT. The cells were washed 3x 5 min in PBS-T and incubated with secondary antibodies conjugated with Alexa fluorophores (1:1000, Invitrogen, A10036, A21208, and A21448) diluted in blocking buffer for 60 min at RT. The cells were then washed twice for 5 min in PBS-T and counterstained for 10 min with Hoechst 33342 (1 μg/mL in PBS). The cells were washed 3x 5 min in PBS, dipped 3x in deionized water and mounted onto microscope slides using buffered glycerol mounting media 90% glycerol (Sigma-Aldrich), 20mM Tris pH 8.0, 0.5% N-propyl gallate (Sigma-Aldrich).

#### Image processing and quantification

For acquisition of immunocytochemistry images, a confocal microscope Leica TCS SPE was used. Images were optimized for brightness and contrast using Fiji-ImageJ. Statistical analysis was conducted using the commercially available software Prism 7.0 (GraphPad Software).

#### qPCR

Total RNA was extracted from cells using TRIzol reagent (Life Technologies) according to the manufacturer’s protocol. After isolation of the aqueous phase 10 μg of glycogen (Thermo Fisher Scientific) was supplemented before precipitation of the RNA. The pellet was resuspended in nuclease free water and the concentration was determined using a NanoDrop. Synthesis of cDNA was performed using the GoScript Reverse Transcriptase kit (Promega) with random hexamer primers, according to the manufacturer’s protocol. Lastly, samples were heat inactivated for 10 min at 70 °C and diluted in nuclease free water. The qPCR was performed in biological triplicates. For the qPCR reaction, 5 μL SYBR green (Roche), 1 μL 10 mM of both forward and reverse primer (see Table S3 for primer sequences) and 10 μL cDNA was mixed in a 10 μL reaction volume and loaded in a white qPCR plate with optical caps (Thermo Fisher Scientific). Samples were spun down briefly and run on a Mx3005P qPCR machine (Agilent) using the following program: 1 cycle of 10 min at 95 °C, 40 cycles of 20 sec at 95 °C, 20 sec at 52 °C, and 30 sec at 72 °C, followed by a melting curve analysis. The Ct values of all samples were first converted from their intrinsic exponential relation to linear related Xo values, using the Xo method (Thomsen et al., 2010). In brief, the triplicate LV2 X o mean values for a sample was normalized to its mean Xo value for GAPDH. The SD was calculated accounting for the common coefficient of variation both for the housekeeping gene and the individual gene investigated. Values represent the gene expression normalized to the expression of GAPDH for each individual sample + SD. Statistical one-way ANOVA, with Tukey’s multiple comparisons test was performed in GraphPad Prism, comparing the expression for one gene between all different samples.

### QUANTIFICATION AND STATISTICAL ANALYSIS

#### Initial quality control and data analysis

Briefly, the sequenced libraries were mapped to pre-mRNA and filtered using 10X Genomics Cell Ranger pipeline. The sequencing data was demultiplexed by bcl2fast (Illumina) which warped in the Cell Ranger followed by aligning to reference genome (Sscrofa11.1 release-94) by STAR (Dobin et al., 2013). Finally, mRNA molecules were counted by Cell Ranger. The quality of the sequencing libraries was assessed by Cell Ranger, which determined in each sample the sequencing depth cutoff that is required for cells to be included in the downstream analysis. All samples were merged for downstream clustering and data analysis, which was performed in Seurat v2 (Satija, Farrell, Gennert, Schier, & Regev, 2015). Similar numbers of UMIs and genes were observed across all batches (Figure S5A, B). Overview of number of cells, mean reads per cell, median genes per cell, and number of reads and pooled brains in the included scRNA-seq libraries can be found in Table S4.

#### Batch correction

We used decomposeVar from the ‘scran’ R package (RRID: SCR_001905) in order to find a list of variable genes that are used for PCA dimensionality reduction; however, to aid the correspondence of single-cell to single-nucleus for down-stream analysis (such as clustering), we identified and excluded genes whose transcripts were highly abundant in empty droplets Empty droplets were defined as cells with <50 UMIs. If the number transcripts that are found in the empty droplet is above 30% the total number of transcripts for a given gene (Figure S5C), that gene will be removed from the list of variable genes.

The individual datasets were log-normalized and the PCA projected gene expression in the datasets were then batch corrected by FastMNN (Haghverdi et al., 2018), which reduced the distance between cell pairs that are found to be reciprocally nearest neighbors across batches (before batch correction; Figure S5D, after correction; Figure 1C). The merged dataset were visualized by t-distributed stochastic neighbour embedding (t-SNE) (van der Maaten & Hinton, 2008).

#### Unsupervised clustering and cell type identification

Cell types were identified using the Louvain algorithm (Blondel, Guillaume, Lambiotte, & Lefebvre, 2008) with a resolution parameter set at 1.6 for the entire dataset, 1.4 for the IN analysis, 1.2 for the oligodendrocyte analysis, and 1.0 for the IP and excitatory neuron analysis. The canonical markers were used to identify the neurons of the clusters (Figure 1D). In addition, we used two reference datasets which each contained smart-seq2 single-cells from human embryonic prefrontal cortex (Fan et al., 2018; Zhong et al., 2018) or from human adult middle temporal gyrus (Astick & Vanderhaeghen, 2018). We used scmap v1.12.0 (Kiselev et al., 2018) to project individual cells onto curated cell-type clusters that are available in each reference. Each cell-type prediction utilizes the consensus of three similarity measures from queried cell to reference cluster centroids using sets of cell-type markers that were identified in the respective reference datasets; however, only the human genes that possess an ortholog gene in pig were used as cell type marker for similarity measure calculation.

#### Differential gene analysis

Marker genes were identified using scfind v3.7.0 (PMID 33649586) based on highest F1 score for cluster specific genes. In principle by: For cluster cj, gene gi is considered a true positive (TP) if it is expressed, a false negative (FN) if it is not expressed, a false positive (FP) if is expressed in a cell assigned to another cluster, and a true negative (TN) if it is not expressed in a cell assigned to a different cluster. For each gi we evaluate precision = TP/(TP+FP), recall = TP/(TP+FN), and F1 = 2*precision*recall/(precision + recall). For each cj, genes are ranked by F1 with the highest scoring genes used as markers.

#### Identification of cell type specific transcription factors

Four excitatory neuron clusters expressing *RELN*, determined as potential SC populations, were investigated for cluster-specific enriched TFs. Porcine genes were defined as TFs (total of 2475 genes) if they, or orthologues, were present in the TFcheckpoint, a list of 3479 specific DNA-binding RNA polymerase II TFs curated by Chawla and colleagues (Chawla et al., 2013). Differentially expressed TFs between each *RELN* cluster and all remaining cells were identified using FindMarkers in the Seurat pipeline (Butler, Hoffman, Smibert, Papalexi, & Satija, 2018), which uses Wilcoxon Rank Sum test. For each of the 2 *RELN* clusters, 20 TFs were selected with the highest log fold-change of average expression between the cluster and the remaining cells.

#### Cell cycle score analysis

Cell cycle phases were scored using the built-in function in Seurat CellCycleScoring, which was used to determine whether a given cell was likely to be in either S, G2M or G1 (which is indistinguishable from G0) phase of cell cycle. Cell cycle scores were based on a list of cell cycle phase-specific genes proposed by Tirosh *et al*. (Tirosh et al., 2016).

## Code availability

All R code used for the analysis of this study is publically available at github: https://github.com/BDD-lab/EC-scRNAseq-2021

## Abbreviations

BMP: bone morphogenetic protein
CHIR: GSK-3α/β inhibitor
CNS: central nervous system
D: Day
DMEM: dulbecco’s modified eagle’s medium
E: Embryonic day
EC: entorhinal cortex
EDTA: Ethylenediaminetetraacetic acid
gDNA: genomic DNA
GOI: gene of interest
h: hours
iPSC: induced pluripotent stem cells
IN: interneurons
IP: intermediate progenitor
L: layer
LV: lentivirus
LEC: lateral entorhinal cortex
MEC: medial entorhinal cortex
MOI: multiplicity of infection
OPC: oligodendrocyte progenitor cell
P: postnatal day
PBS: phosphate-buffered saline
PCR: polymerase chain reaction
PYR: pyramidal neuron
qPCR: quantitative real time PCR
*RELN*: Reelin gene
RT: room temperature
scRNA-seq: Single-cell RNA sequencing
SC: stellate cell
SD: standard deviation
SPE: speed cell
SST: somatostatin
TF: transcription factor
U: units of viral particles
UMI: unique molecular identifier
VPD: volume of produced virus

## Declarations

### Ethics approval and consent to participate

The experiments conform to the relevant regulatory standards for acquisition and use of fetal pig material.

### Consent for publication

All authors consent that this research is suitable for publication.

### Availability of data and materials

The datasets generated are available at the NCBI repository with GEO accession number GSE134482 (https://www.ncbi.nlm.nih.gov/geo/query/acc.cgi?acc=GSE134482). The submitted data includes the raw sequencing data as fastq files together with the processed count matrix used in this study.

### Authors’ contributions

TB: Performed scRNA-seq, direct programming, analyzed data and contributed to the writing of the manuscript

YL: Performed scRNA-seq, analyzed data, designed the GOIs subclone pipeline and contributed to the writing of the manuscript

LM: Performed lentiviral production.

JL: Performed scRNA-seq analysis.

UP: Performed scRNA-seq analysis

L-FH: Contributed to methodology of bioinformatic analysis of the scRNA-seq data

AAM: Performed scRNA-seq analysis

ILV; Performed scRNA-seq analysis

MD: Supplied Ngn2 backbones and Ngn2 lentivirus

SES: Contributed to methodology of bioinformatic analysis of the scRNA-seq data

JTHL: Contributed to methodology of bioinformatic analysis of the scRNA-seq data

NP: Contributed to methodology of bioinformatic analysis of the scRNA-seq data

MP: Contributed to methodology of bioinformatic analysis of the scRNA-seq data

BRK: Contributed to methodlogy of scRNA-seq

MD: Contributed with lentiviral plasmids for direct programming experiments and viral tittering methodology

PH: Supervised project and contributed to acquisition of funding

MPW: Contributed to anatomical analysis of the EC in the pig and in acquisition of funding

JG: Provided server access for scRNA-seq data, supervised the project and contributed to interpretation of bioinformatic data

MH: Provided software, personnel and resources for bioinformatic analyses of the scRNA-seq and contributed to interpretation of the scRNA-seq data

THP: Provided materials, personnel and resources for scRNA-seq, supervised project and contributed to interpretation of the scRNA-seq data

KK: Provided materials, personnel and resources for scRNA-seq, supervised project and contributed to interpretation of the data

VH: Contributed to the project’s concept, methodology, formal analyses, project’s resources, writing, reviewing & editing the manuscript. Administered and managed the project and acquired the project’s funding.

All authors read and approved the final manuscript

## Acknowledgements

We thank Per Torp Sangild for his donation of healthy adult sow brains for this study. We used the Single Cell Genomics Facility at BRIC for access to 10X Chromium platform. We also acknowledge the help from Anna Fossum at the FACS Facility at BRIC, University of Copenhagen for help with the FACS sorting.

---

**Figure S1.**
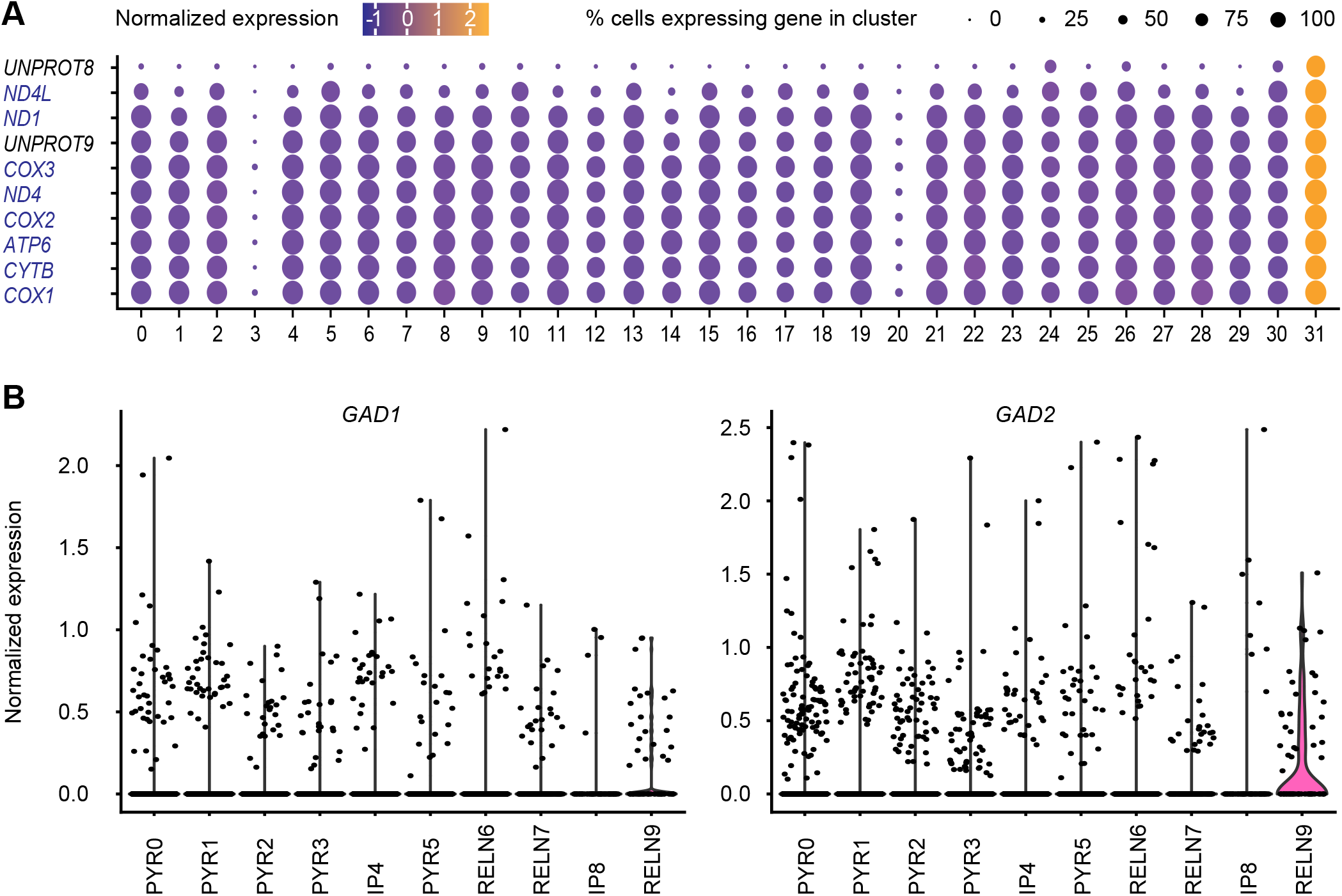
Mitochondrial gene expression in the whole scRNA-seq dataset, evaluation of interneuron (IN) genes in the entorhinal cortex (EC) excitatory neuron/intermediate progenitor (IP) clusters. **(A)** The top ten gene features of cluster 31 identified in Seurat using FindAllMarkers (with Wilcoxon Rank Sum test) showed enrichment of mitochondrial genes in the cluster, likely representing dying cells, which resulted in the cluster being removed from further analyses. **(B)** Expression of the IN markers, GAD1 and GAD2 in the IP and excitatory neuron clusters highlight low expression across all the subtypes, with the exception of GAD2 in PYR12 which suggests it may be an IN progenitor.

**Figure S2.**
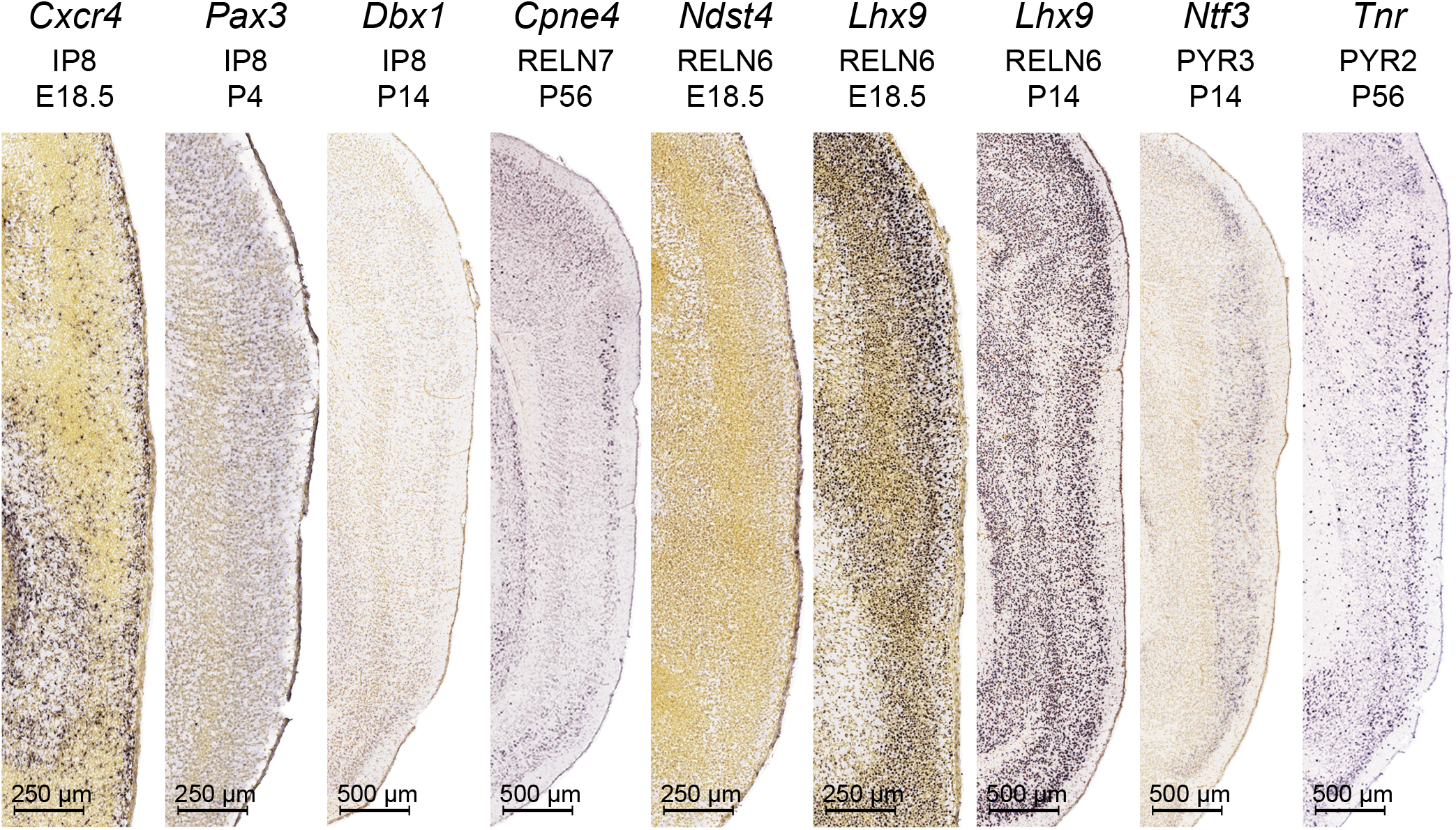
Expression of RELN, PYR and IP cluster markers in saggital sections of the entorhinal cortex (EC) from the Allen Developing Mouse Brain Atlas and Allen Mouse Brain Atlas. Expression of select markers for varying excitatory neuron clusters highlighted in Figure 4C were investigated in ISH sagittal sections revealing putative locations of the cell types, such as RELN7 and PYR2 neurons residing in the superficial layers. Image credit: Allen Institute (82-84). Scale bars: 250 μm (for E18.5 and P4) and 500 μm (for P14 and P56).

**Figure S3.**
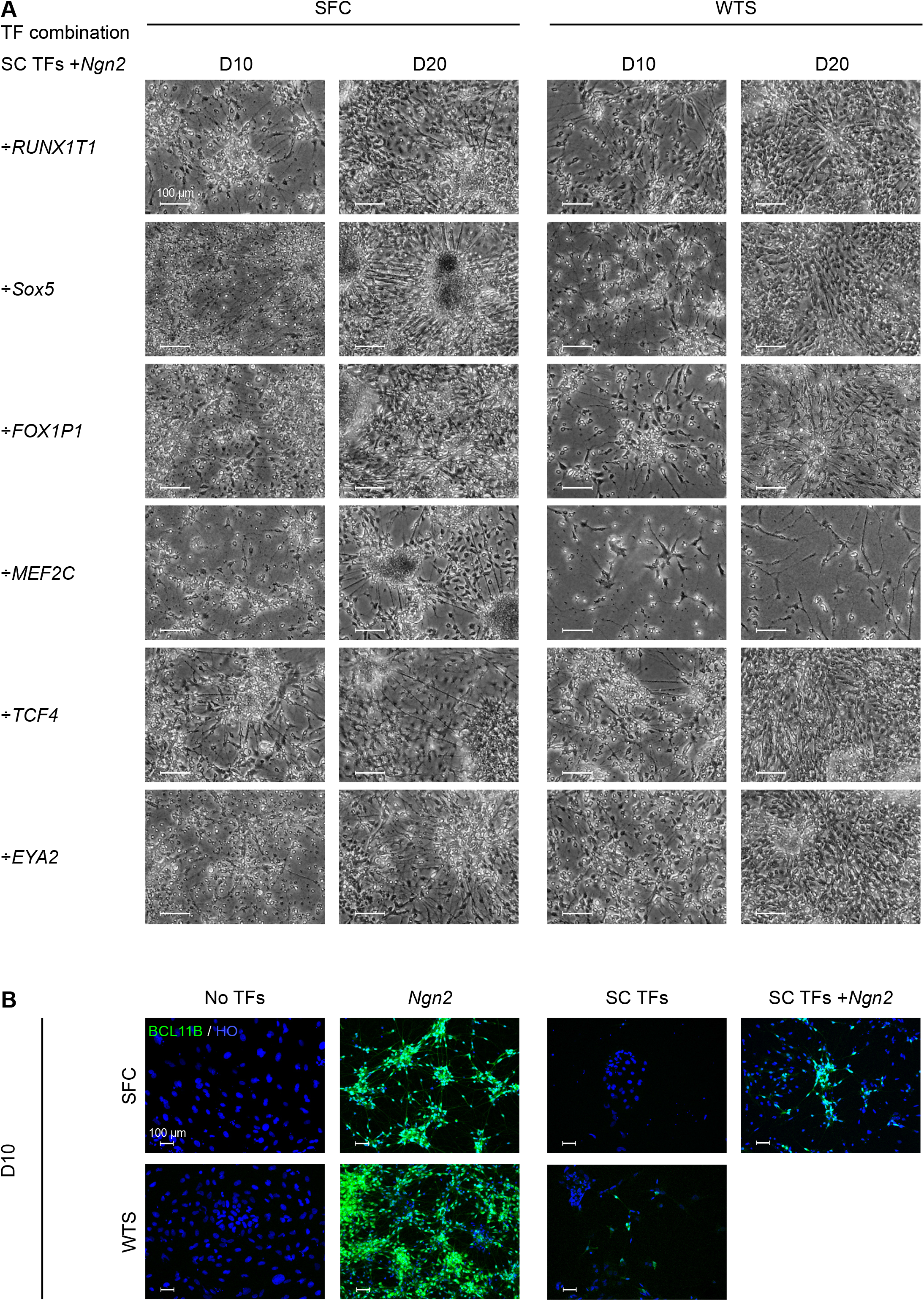
Direct programming of induced pluripotent stem cells (iPSCs) using a leave-one-out approach of the stellate cell-specific transcription factors in the presence of Ngn2 resulted in the production of stellate cell-like cells. **(A)** Morphological changes of the two iPSC lines, SFC 180-01-01 (SFC) and WTSli024A (WTS) following transduction with different combinations of five out of the six stellate cell transcription factors (SC TFs), together with Ngn2 at day (D)10 and D20. Scale bar: 100 μm. **(B)** Expression of BCL11B in the SFC and WTS cell lines at D10 after transduction with the SC TFs alone, the SC TFs with Ngn2, Ngn2 alone, or with no TFs. HO; Hoechst. Scale bar: 100 μm.

**Figure S4.**
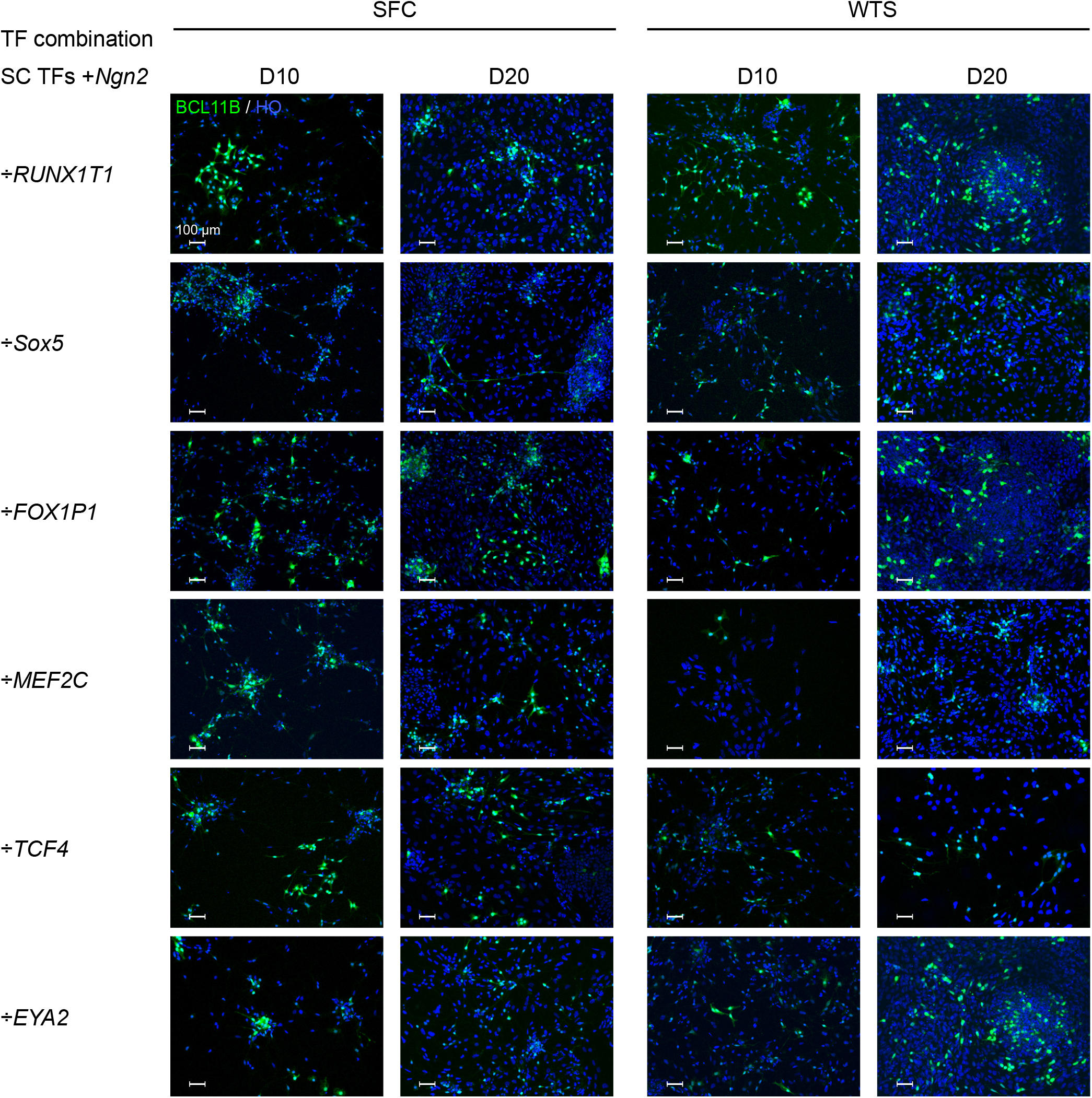
BCL11B expression in induced pluripotent stem cells (iPSCs) following different combinations of induced transcription factor expression. BCL11B expression was observed in both iPSC lines SFC 180-01-01 (SFC) and WTSli024A (WTS) transduced with different combinations of five stellate cell transcription factors (SC TFs) in the presence of Ngn2, at Day (D)10 and 20 post transduction. HO; Hoechst. Scale bar: 100 μm.

**Figure S5.**
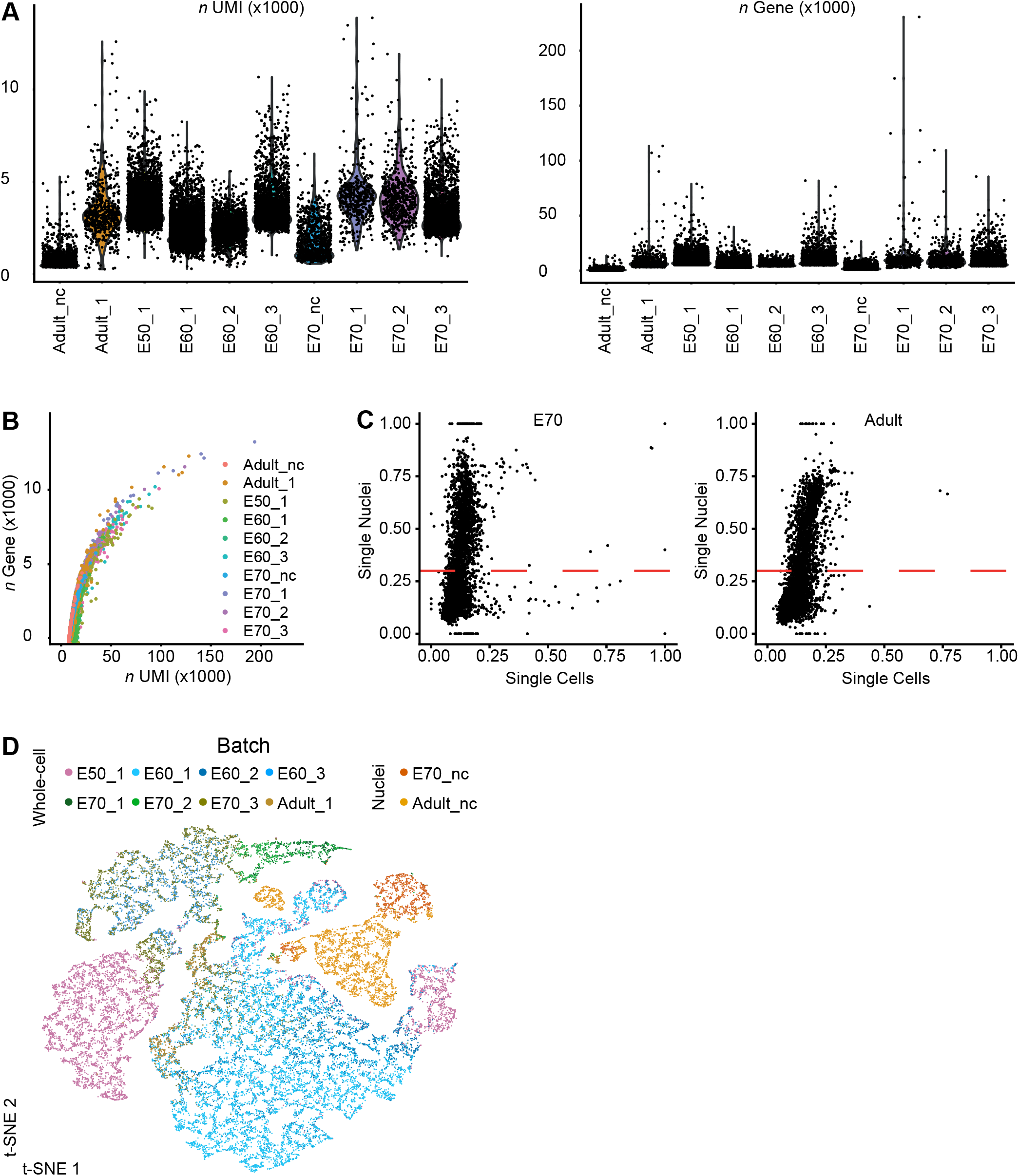
Validation data for the scRNA-seq experiments. **(A)** Violin plots of the number (n) of UMIs and genes in each single-cell sequencing batch. **(B)** Scatter plots of the number (n) of UMIs versus genes for each single-cell sequencing batch. **(C)** Fraction of transcripts out of total transcripts for a given gene in empty droplets at representative stages (E70 and Adult), for both isolated nuclei (nc) and whole cell samples. Genes with a fraction above 0.30 were not included as variable genes in the downstream analysis. Abbreviations: n, number; E, embryonic **(D)** A t-SNE plot of the scRNA-seq dataset prior to fastMNN batch correction.

**Table S1:** Human ortholog names identified from pig ensemble names.

| **Ensemble Name** | **Human ortholog names** |
| --- | --- |
| ENSSSCG00000010212 | ANK3 |
| ENSSSCG00000011178 | CPNE4 |
| ENSSSCG00000015545 | GLUL |
| ENSSSCG00000009327 | HMGB1 |
| ENSSSCG00000011267 | MOBP |
| ENSSSCG00000015401 | PCLO |
| ENSSSCG00000007644 | PILRB |
| ENSSSCG00000016219 | RESP18 |
| ENSSSCG00000029160 | UNPROT1 |
| ENSSSCG00000006963 | UNPROT2 |
| ENSSSCG00000032935 | UNPROT3 |
| ENSSSCG00000017019 | UNPROT5 |
| ENSSSCG00000034554 | UNPROT8 |
| ENSSSCG00000035520 | UNPROT9 |
| ENSSSCG00000036465 | USP6NL |
| ENSSSCG00000005658 | ZDHHCL2 |
| ENSSSCG00000005063 | OTX2 |
| ENSSSCG00000027279 | PTHR |

**Table S2:**
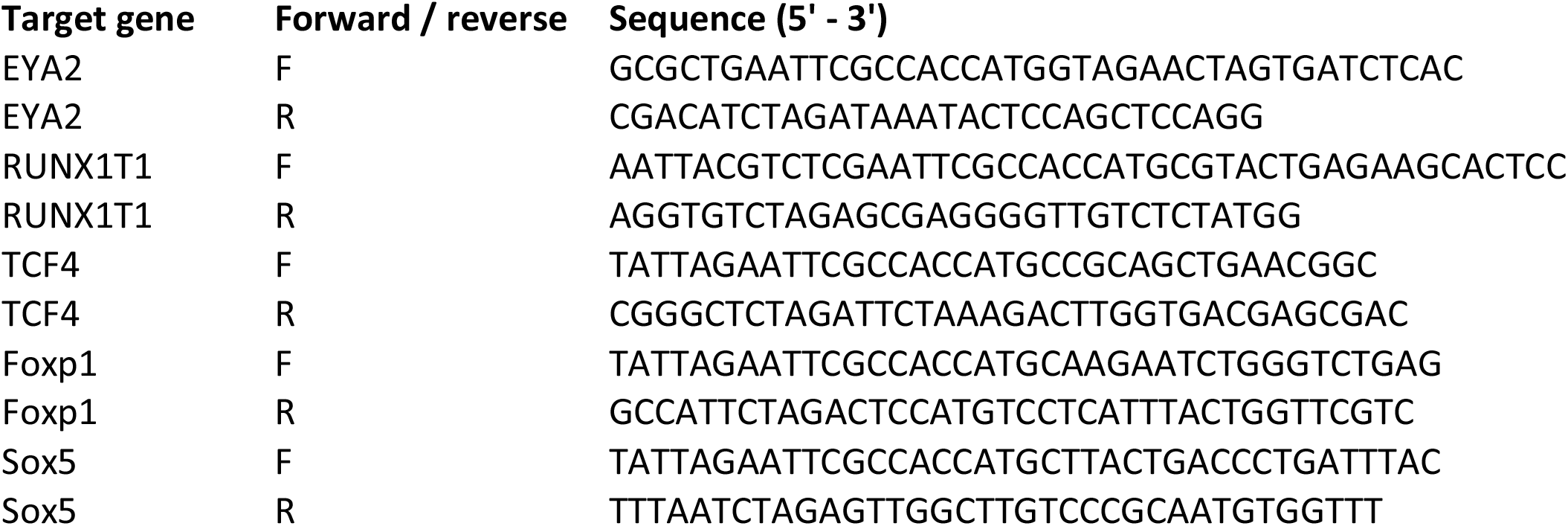
Primers for cloning of plasmids.

**Table S3:**
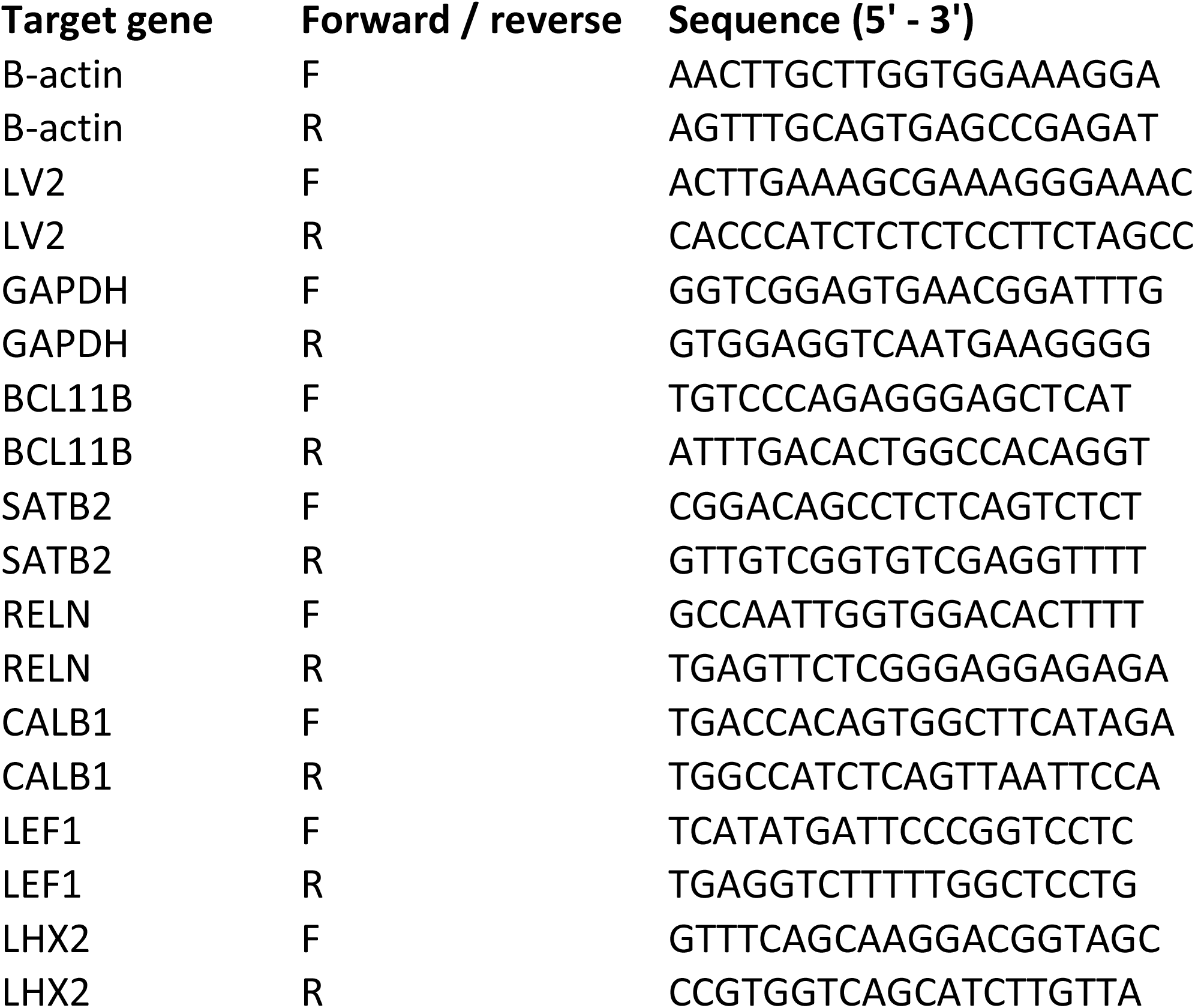
Primers for qPRC.

**Table S4:**
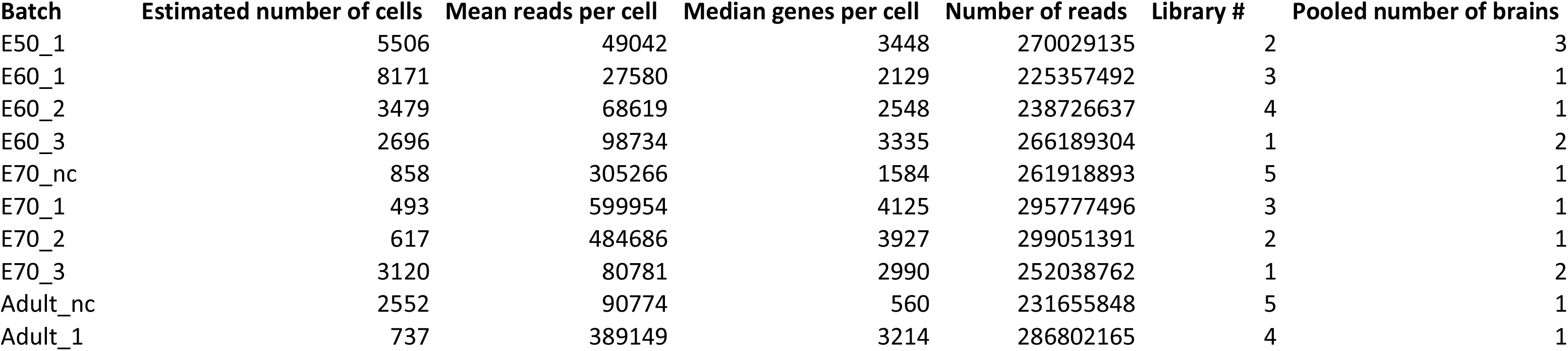
Overview of batches, captured cells, mean reads, number of libraries, and number of brains used for scRNA-seq
experiments.

| **Batch** | **Estimated number of cells** | **Mean reads per cell** | **Median genes per cell** | **Number of reads** | **Library #** | **Pooled number of brains** |
| --- | --- | --- | --- | --- | --- | --- |
| E50_1 | 5506 | 49042 | 3448 | 270029135 | 2 | 3 |
| E60_1 | 8171 | 27580 | 2129 | 225357492 | 3 | 1 |
| E60_2 | 3479 | 68619 | 2548 | 238726637 | 4 | 1 |
| E60_3 | 2696 | 98734 | 3335 | 266189304 | 1 | 2 |
| E70_nc | 858 | 305266 | 1584 | 261918893 | 5 | 1 |
| E70_1 | 493 | 599954 | 4125 | 295777496 | 3 | 1 |
| E70_2 | 617 | 484686 | 3927 | 299051391 | 2 | 1 |
| E70_3 | 3120 | 80781 | 2990 | 252038762 | 1 | 2 |
| Adult_nc | 2552 | 90774 | 560 | 231655848 | 5 | 1 |
| Adult_1 | 737 | 389149 | 3214 | 286802165 | 4 | 1 |

